# Linker histone H1-0 is a specific mediator of the repressive ETV6::RUNX1 transcriptional landscape

**DOI:** 10.1101/2024.06.28.601221

**Authors:** Vera H. Jepsen, Andrea Hanel, Daniel Picard, Juha Mehtonen, Rebecca Hasselmann, Julian Schliehe-Diecks, Katerina Scharov, Jia-Wey Tu, Rigveda Bhave, Ersen Kameri, Nan Qin, Herui Wang, Zhengping Zhuang, Rabea Wagener, Lena Blümel, Tobias Lautwein, Daniel Hein, Gesine Kögler, Marc Remke, Sanil Bhatia, Merja Heinäniemi, Arndt Borkhardt, Ute Fischer

## Abstract

*ETV6::RUNX1* is the most common oncogenic fusion in pediatric B cell precursor acute lymphoblastic leukemia (BCP-ALL). It induces a clinically silent preleukemic state that requires secondary mutations for progression to leukemia. However, the molecular mechanisms contributing to the characteristic quiescence of *ETV6::RUNX1*+ preleukemic cells remain elusive. Here, we detect factors involved in the preleukemic state by generating human induced pluripotent stem cell (hiPSC) models using CRISPR/Cas9 gene editing. We identified upregulation of linker histone *H1-0* in our preleukemic models, which was preserved upon hematopoietic differentiation and transformation to BCP-ALL. ETV6::RUNX1 induces *H1-0* promoter activity whereas depletion of H1-0 specifically inhibited ETV6::RUNX1 signature genes, indicating its role as a key mediator of the ETV6::RUNX1 transcriptome. Single-cell gene expression analysis revealed high *H1-0* levels in quiescent cells during hematopoiesis and inverse correlation with transcriptional activity. Pharmacologically, H1-0 protein levels correspond to susceptibility of BCP-ALL towards histone deacetylase inhibitors (HDACi). Altogether, our study provides novel insights into ETV6::RUNX1-induced quiescence and suggests that further investigation into combinatorial treatment of BCP-ALL using the H1-0- inducing HDACi Quisinostat may be worthwhile.

## Introduction

The chromosomal translocation t(12;21)(p13;q22) is the most common structural variation of pediatric B cell precursor acute lymphoblastic leukemia (BCP-ALL) and results in the fusion of the two hematopoietic transcription factors ETS translocation variant 6 (*ETV6*) and runt-related transcription factor 1 (*RUNX1*). The *ETV6::RUNX1* fusion gene is acquired *in utero* in 1-5% of newborns (Mori et al, 2002; Schäfer et al, 2018) and requires further oncogenic mutations for progression to overt leukemia, predominantly including copy number alterations of genes involved in B cell development or cell cycle, e.g. *ETV6*, *PAX5*, *CDKN2A* and *CDKN2B* (Brady et al, 2022; Greaves, 2018; Mullighan et al, 2007; Papaemmanuil et al, 2014). Despite overall survival rates of *ETV6::RUNX1*+ pediatric leukemia exceeding 90% with current chemotherapy regimens, patients suffer from substantial acute and late toxicities. Disease recurrence, particularly late-onset relapses, is observed in approximately 20% of patients (Gu et al, 2019; Konrad et al, 2003). This underlines the importance of understanding *ETV6::RUNX1*+ BCP-ALL pathophysiology to enable further improvement of treatment.

While ETV6::RUNX1 itself is not sufficient for leukemic transformation, we and others have demonstrated that the fusion protein establishes a distinct preleukemic cell state (Böiers et al, 2018; Linka et al, 2014; Linka et al, 2013; Wray et al, 2022). ETV6::RUNX1 exerts an overall repressive effect on preleukemic cells, impeding early B cell differentiation, cell cycle and inflammatory pathways, such as TGFβ signaling (Böiers et al, 2018; Ford et al, 2009; Teppo et al, 2016; Wray et al, 2022). Transcriptional repression is conferred via the pointed domain (PNT) of the ETV6 moiety, while the runt-homology domain (RHD) of the RUNX1 fusion part directly binds to promoters harboring the canonical RUNX1-binding motif ‘TGYGGTY’ (Fenrick et al, 1999; Hiebert et al, 1996b; Morrow et al, 2007; Zelent et al, 2004). ETV6::RUNX1 associates with multiple co-repressors, including NCOR1, mSin3A and histone deacetylases, such as HDAC3 (Guidez et al, 2000; Wang & Hiebert, 2001), that induce changes in chromatin structure, leading to the characteristic repression of RUNX1 target genes (Starkova et al, 2007).

Modeling approaches of *ETV6::RUNX1*+ preleukemia and overt leukemia in mice were largely unable to reproduce restriction to B lineage leukemia seen in humans (Bernardin et al, 2002; Schindler et al, 2009; van der Weyden et al, 2011). This might be attributed to expression level-dependent effects of ETV6::RUNX1, especially in models using viral transduction (Tsuzuki & Seto, 2013). Additionally, discrepancies between *ETV6::RUNX1*+ mouse and human models were linked to poor inter-species conservation of GGAA repeat enhancers recently identified as key regulators of the *ETV6::RUNX1*+ BCP-ALL gene signature (Kodgule et al, 2023). Therefore, accurately recapitulating the intricate effects of ETV6::RUNX1 may necessitate modeling its function in a human background with physiological expression levels, as demonstrated using preleukemic *ETV6::RUNX1*+ human induced pluripotent stem cells (hiPSCs) that exhibit a transient block of early B lymphopoiesis *in vitro* (Böiers et al, 2018).

In this study, we detect consistent upregulation of linker histone H1-0 in a preleukemic hiPSC model and leukemic blasts carrying the *ETV6::RUNX1* fusion gene. As a member of the H1 family of linker histones, H1-0 affects chromatin compaction (Morales Torres et al, 2016; Willcockson et al, 2021). H1-0 is heterogeneously expressed in solid tumors where it contributes to the intricate balance between cancer cell proliferation and differentiation (Morales Torres et al, 2016). Our data reveals that H1-0 acts as an important mediator of the ETV6::RUNX1 gene expression profile and identifies the HDACi Quisinostat as a potential targeted approach for combinatorial drug treatment of *ETV6::RUNX1*+ leukemic cells.

## Results

### ETV6::RUNX1 induces HSC expansion and decreased transcriptional activity

To analyze specific gene expression patterns of *ETV6::RUNX1*-translocated preleukemia in models without additional secondary alterations, we generated monoclonal hiPSC lines derived from two donors (termed HW8 and ChiPSC12). We used a CRISPR/Cas9-mediated knock-in approach to directly fuse *RUNX1* exons 2-8 to *ETV6* exon 5 and to place the resulting fusion gene under the physiological control of the endogenous *ETV6* promotor (Fig. 1A). We confirmed correct sequence of the *RUNX1* insert by genotyping polymerase chain reaction (PCR) of the *ETV6* locus (Fig. EV1A-B) and Sanger sequencing (Fig. EV1C). *ETV6::RUNX1* levels in the hiPSC lines as detected by reverse transcription quantitative PCR (RT-qPCR) and Western blot were lower compared to the *ETV6::RUNX1*+ BCP-ALL cell line REH (Figs. EV1D and 1B). All hiPSC lines maintained typical hiPSC microscopic morphology and expression of the pluripotency markers *SSEA-4*, *DNMT3B*, *GDF3*, *POU5F1* and *NANOG* as determined by flow cytometric analyses and RT-qPCR; chromosomal integrity was confirmed by karyotype analysis (Fig. EV2). All *ETV6::RUNX1*+ hiPSC lines harbored a monoallelic insertion of the *RUNX1* HDR template at the *ETV6* locus as detected by PCR and Sanger sequencing (Fig. EV1E-G). Since the *RUNX1* HDR template disrupts one *ETV6* allele, expression of full-length *ETV6* was lower in the CRISPR/Cas9-edited hiPSCs compared to the wild-type controls (Fig. 1B-C). This aligns with the genetic profile of *ETV6::RUNX1*+ ALL patients who commonly exhibit heterozygosity for the fusion gene. Since *ETV6* exon 6 is not retained in the *ETV6::RUNX1* fusion gene, full-length *ETV6* was detected using an RT-qPCR spanning exons 5 and 6 (Fig. EV1H). REH cells served as negative control due to deletion of the remaining copy of *ETV6* (Fig. 1B-D).

**Figure 1.**
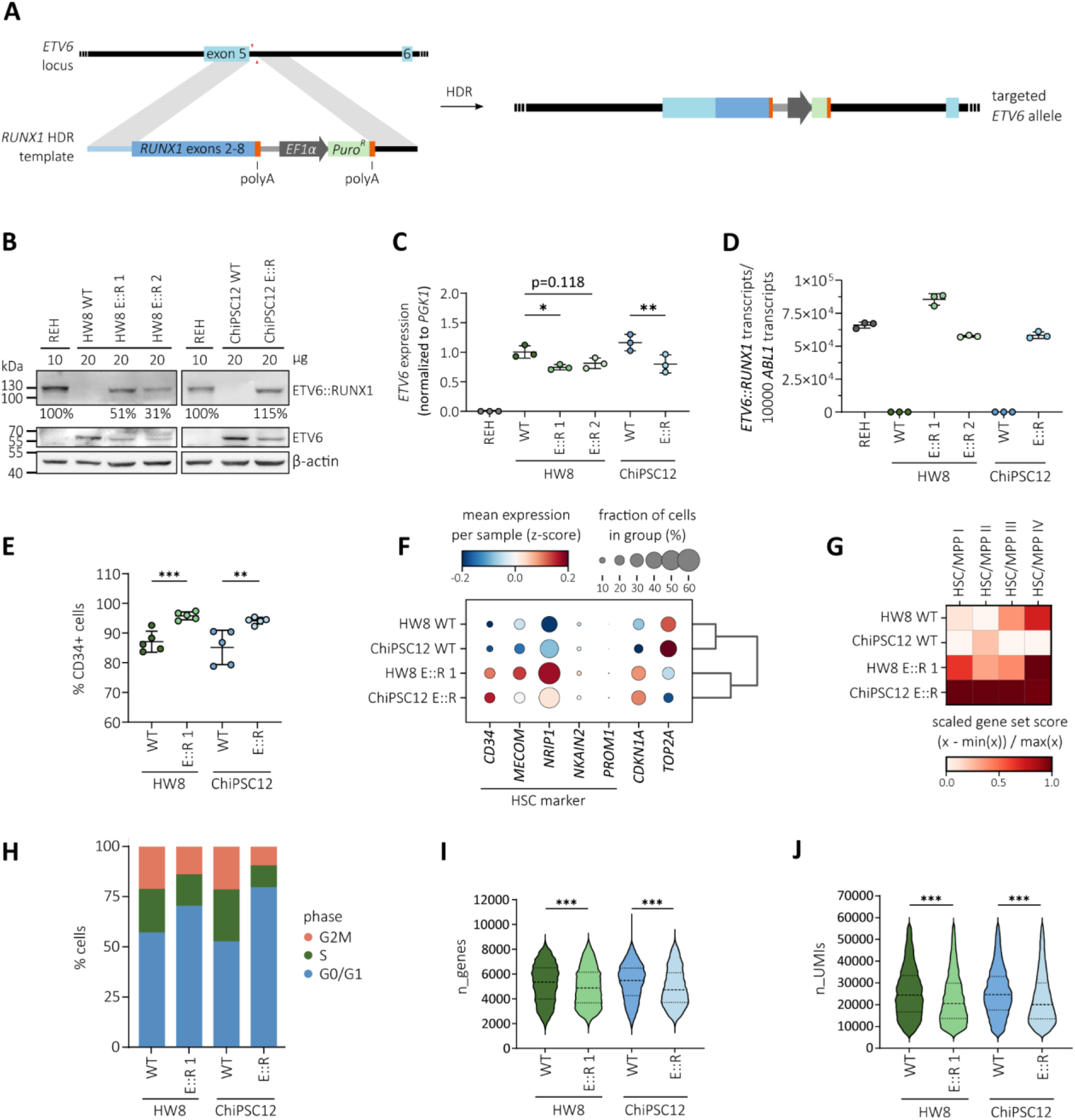
ETV6::RUNX1 induces HSC expansion and decreased transcriptional activity. **(A)** Scheme representing CRISPR/Cas9-mediated targeted editing of the endogenous *ETV6* locus using a *RUNX1* homology-directed repair (HDR) template. **(B)** Western blot analysis of REH and hiPSC lysates using antibodies directed against β-actin, ETV6 and RUNX1 detecting the ETV6::RUNX1 (E::R) fusion protein. **(C)** Relative *ETV6* expression determined by RT-qPCR in REH and hiPSCs. Mean expression ± standard deviation is indicated and data was analyzed for statistical significance using an ordinary one-way ANOVA (*p<0.05, **p<0.01). **(D)** Quantification of *ETV6::RUNX1* expression by RT-qPCR in REH and hiPSC-derived HPCs. Mean expression ± standard deviation is indicated. **(E)** Frequencies of CD34+ hiPSC-derived HPCs determined by flow cytometry. Each dot represents a technical replicate (n=5). Replicates were pooled for scRNA-seq. Mean expression ± standard deviation is indicated. Data was analyzed for statistical significance using an unpaired t-test (**p<0.01, ***p<0.001). **(F)** Dot plot indicating mean expression per sample (z-score) of HSC marker genes *CD34*, *MECOM*, *NRIP1*, *NKAIN2* and *PROM1*, cell-cycle inhibitor *CDKN1A* (p21) and G2/M marker DNA topoisomerase *TOP2A* determined by scRNA-seq. Gene expression frequency (fraction of cells per sample) is indicated by dot size and log-transformed, normalized and scaled (z-score) gene expression level is indicated by color intensity. **(G)** Heat map visualizing scaled HSC/MPP I-IV scores of scRNA-seq data derived from *ETV6::RUNX1*+ and WT HPCs using gene set signatures derived from Jardine *et al*. (https://developmental.cellatlas.io/fetal-bone-marrow, accession number E-MTAB-9389 (Jardine et al, 2021)). **(H)** Stacked bar plot showing distribution of cell cycle stages of *ETV6::RUNX1*+ and WT HPCs analyzed by scRNA-seq. **(I, J)** Violin plots depicting number of expressed genes (n_genes) and unique molecular identifier counts (n_UMIs) per cell detected by scRNA-seq. Median expression and quartiles are indicated. Data was analyzed for statistical significance using an unpaired t-test (***p<0.001).

To delineate effects of *ETV6::RUNX1* in hematopoietic cells, we performed scRNA-seq analysis of hematopoietic progenitor cells (HPCs) differentiated from *ETV6::RUNX1*+ and wild-type hiPSCs. As confirmed by RT-qPCR, *ETV6::RUNX1* expression in HPCs increased to levels comparable with REH cells (Fig. 1D). After 12 days of differentiation, we observed significantly increased numbers of CD34+ cells from *ETV6::RUNX1*+ hiPSCs compared to wild-type hiPSCs (Fig. 1E). Correspondingly, scRNA-seq identified upregulation of hematopoietic stem cell (HSC) marker genes *CD34*, *MECOM*, *NRIP1*, *NKAIN2* and *PROM1* (Heumos et al, 2023), and enrichment of HSC/multipotent progenitor (MPP) signatures (Jardine et al, 2021) in *ETV6::RUNX1*+ HPCs (Fig. 1F-G). Furthermore, cell cycle scoring revealed a skewed distribution towards the G0/G1 cell cycle phase in *ETV6::RUNX1*+ HPCs (Fig. 1H), reflected by increase of cell cycle kinase inhibitor *CDKN1A* expression and decrease of G2/M phase-specific DNA topoisomerase *TOP2A* (Fig. 1F). Overall, transcriptional diversity and activity was reduced in *ETV6::RUNX1*+ HPCs as indicated by lower number of expressed genes per cell and unique transcripts detected per cell, respectively (Fig. 1I-J).

Altogether, our preleukemic hiPSC model recapitulates the repressive effect of the *ETV6::RUNX1* fusion gene (Fuka et al, 2011a; Teppo et al, 2016; Wray et al, 2022), inducing accumulation of HSCs upon hematopoietic differentiation and increasing quiescence *in vitro*.

### *H1-0* is consistently upregulated in preleukemia and BCP-ALL expressing *ETV6::RUNX1*

To identify potential genes that contribute to reduced transcriptional diversity and activity detected in *ETV6::RUNX1*+ preleukemia, we performed bulk RNA-seq of *ETV6::RUNX1*+ and wild-type hiPSCs. Principal component analysis (PCA) clearly separated samples according to genotype (Fig. 2A). Altogether, we found consistent differential expression of 20 genes with an absolute fold change >2 and p<0.05 in the three *ETV6::RUNX1*+ hiPSC lines (Figs. 2B and EV3, Datasets EV1-3). Among these genes, *H1-0* has previously been identified as the most significantly upregulated gene in dormant leukemia stem cell (LSC)-like cells (Ebinger et al, 2016). As a linker histone, H1-0 is involved in epigenetic regulation of chromatin and affects cellular differentiation states (Morales Torres et al, 2016), making it a compelling candidate for further investigation. Elevated levels of *H1-0* identified by RNA-seq in the *ETV6::RUNX1*+ hiPSCs were confirmed both by RT-qPCR (2.4-fold increased mean expression; Fig. 2C) and Western blot (Fig. 2D). Moreover, upregulation of *H1-0* in *ETV6::RUNX1*+ preleukemic cells is preserved during differentiation of hiPSCs along the B lymphoid lineage (Fig. 2E; data is derived from a published RNA-seq dataset, accession number E-MTAB-6382 (Böiers et al, 2018)).

**Figure 2.**
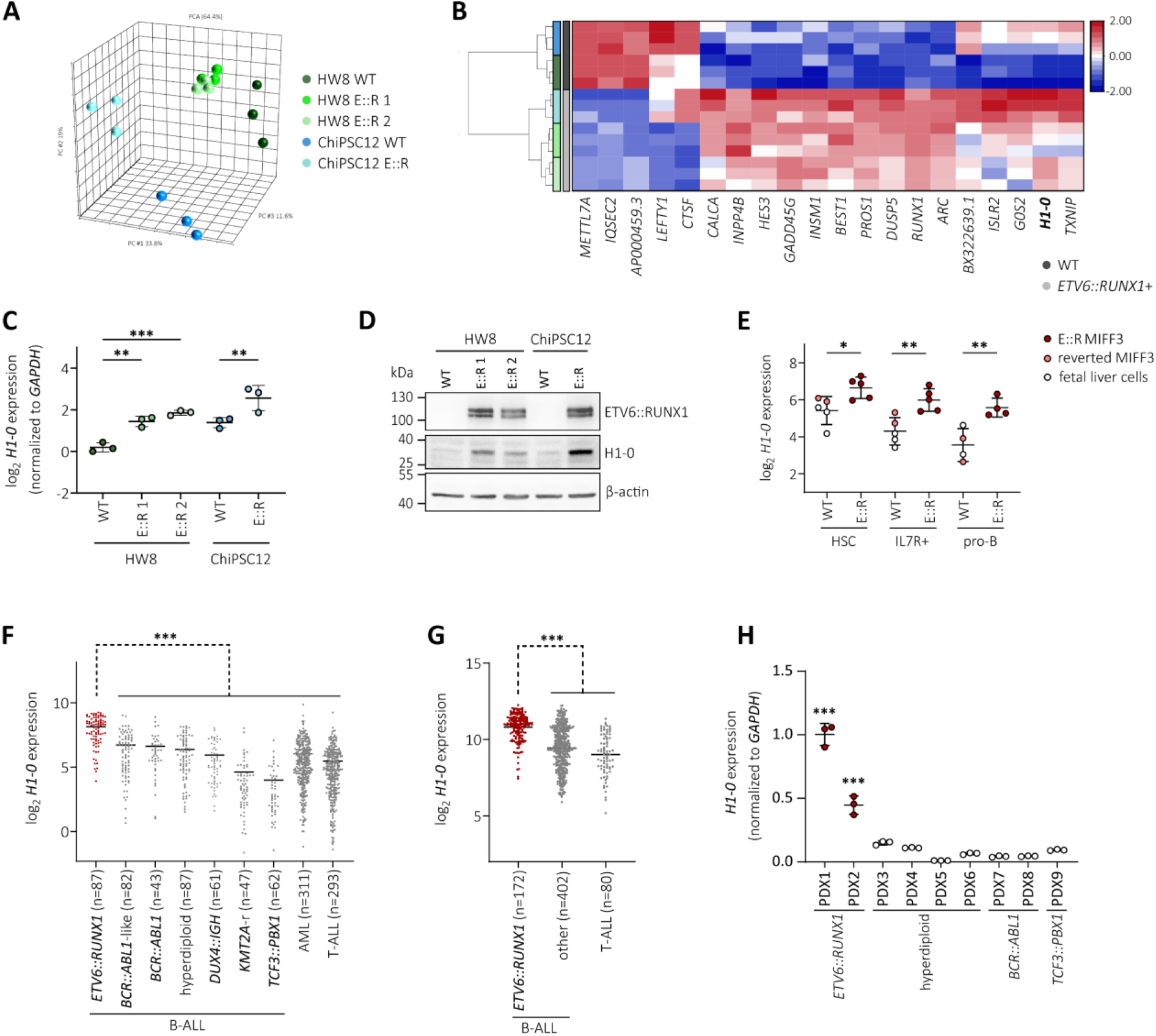
*H1-0* is consistently upregulated in preleukemia and BCP-ALL expressing *ETV6::RUNX1*. **(A)** Principal component analysis (PCA) plot of *ETV6::RUNX1*+ (E::R) and WT hiPSC transcriptome profiles based on all detected genes (n=16328). **(B)** Hierarchical clustering analysis of differentially expressed genes (absolute fold change >2 and p<0.05) between *ETV6::RUNX1*+ and wild-type (WT) hiPSCs detected by RNA-seq. **(C)** *H1-0* expression levels determined by RT-qPCR in *ETV6::RUNX1*+ and WT hiPSCs subjected to RNA-seq. Values were normalized to HW8 WT expression levels as well as to *GAPDH* expression. **(D)** Representative Western blot analysis of ETV6::RUNX1, H1-0, ETV6 and β-actin levels in *ETV6::RUNX1*+ and WT hiPSCs. **(E)** *H1-0* levels in HSCs (CD19-CD34+CD45RA-), IL7R+ (CD19-CD34+CD45RA+IL7R+) and pro-B (CD19+CD34+) cells differentiated from *ETV6::RUNX1*+ or reverted MIFF3 hiPSCs, and fetal liver cells. Data is derived from an RNA-seq dataset by Böiers *et al*. (accession number E-MTAB-6382 (Böiers et al, 2018)). Data was analyzed for statistical significance using an ordinary one-way ANOVA (*p<0.05, **p<0.01). **(F, G)** *H1-0* levels across two leukemia patient cohorts derived from the (F) PeCan St. Jude database (Downing et al, 2012; McLeod et al, 2021) and (G) an expression microarray dataset (accession number GSE87070 (Polak et al, 2019)). Number of patients per leukemia entity and mean expression is indicated. Data was analyzed for statistical significance using an ordinary one-way ANOVA (***p<0.001). **(H)** *H1-0* expression was quantified by RT-qPCR in patient-derived xenograft (PDX) samples (n=9). Mean expression ± standard deviation is shown.

We previously found *H1-0* expression to be restricted to *ETV6::RUNX1*+ bone marrow blasts compared to peripheral blood CD19+ cells (Linka et al, 2013), indicating that *H1-0* upregulation is preserved upon leukemic transformation and highly specific for leukemic cells carrying the *ETV6::RUNX1* fusion gene. To confirm this finding, we analyzed transcriptomic data derived from two patient cohorts encompassing a total of 1,727 leukemia patient samples (PeCan St. Jude cohort (Downing et al, 2012; McLeod et al, 2021) and GSE87070 (Polak et al, 2019)). Additionally, we determined *H1-0* expression in nine patient-derived xenograft (PDX) samples by RT-qPCR (n=2 *ETV6::RUNX1*+, n=4 high-hyperdiploid, n=2 *BCR::ABL1*+, n=1 *TCF3::PBX1*+ BCP-ALL). Across all cohorts, *ETV6::RUNX1*+ BCP-ALL showed significantly elevated *H1-0* levels compared to other leukemia entities (Figs. 2F-H).

### *ETV6::RUNX1* expression is sufficient to induce *H1-0* promoter activation

Given that our findings show strong association between *ETV6::RUNX1* and *H1-0* expression, we tested the potential of ETV6::RUNX1 to transactivate the *H1-0* promoter. To this end, we cloned the *H1-0* promoter region (−351 to +161 from TSS) into a luciferase reporter plasmid (Fig. 3A), which was transfected into 293T cells along with either an empty vector or vectors containing FLAG-tagged *ETV6::RUNX1* or *RUNX1* sequences. Luciferase activity measurements confirmed that expression of *ETV6::RUNX1* is sufficient to activate the *H1-0* promoter (2.2-fold), while *RUNX1* expression induces reduction of luciferase activity (3.1-fold; Fig. 3B). However, our previous analyses in murine cells (Linka et al, 2014; Linka et al, 2013) and analysis of the *H1-0* promoter region using published chromatin immunoprecipitation sequencing (ChIP-seq) data of *ETV6::RUNX1*+ REH cells (accession numbers GSE176084 (Jakobczyk et al, 2022) and GSE117684 (Jakobczyk et al, 2021)) did not show direct binding of either the fusion protein or RUNX1 to the *H1-0* promoter region or distal enhancer regions upstream of *H1-0* (Fig. EV4), suggesting an indirect mechanism of *H1-0* upregulation upon *ETV6::RUNX1* expression.

**Figure 3.**
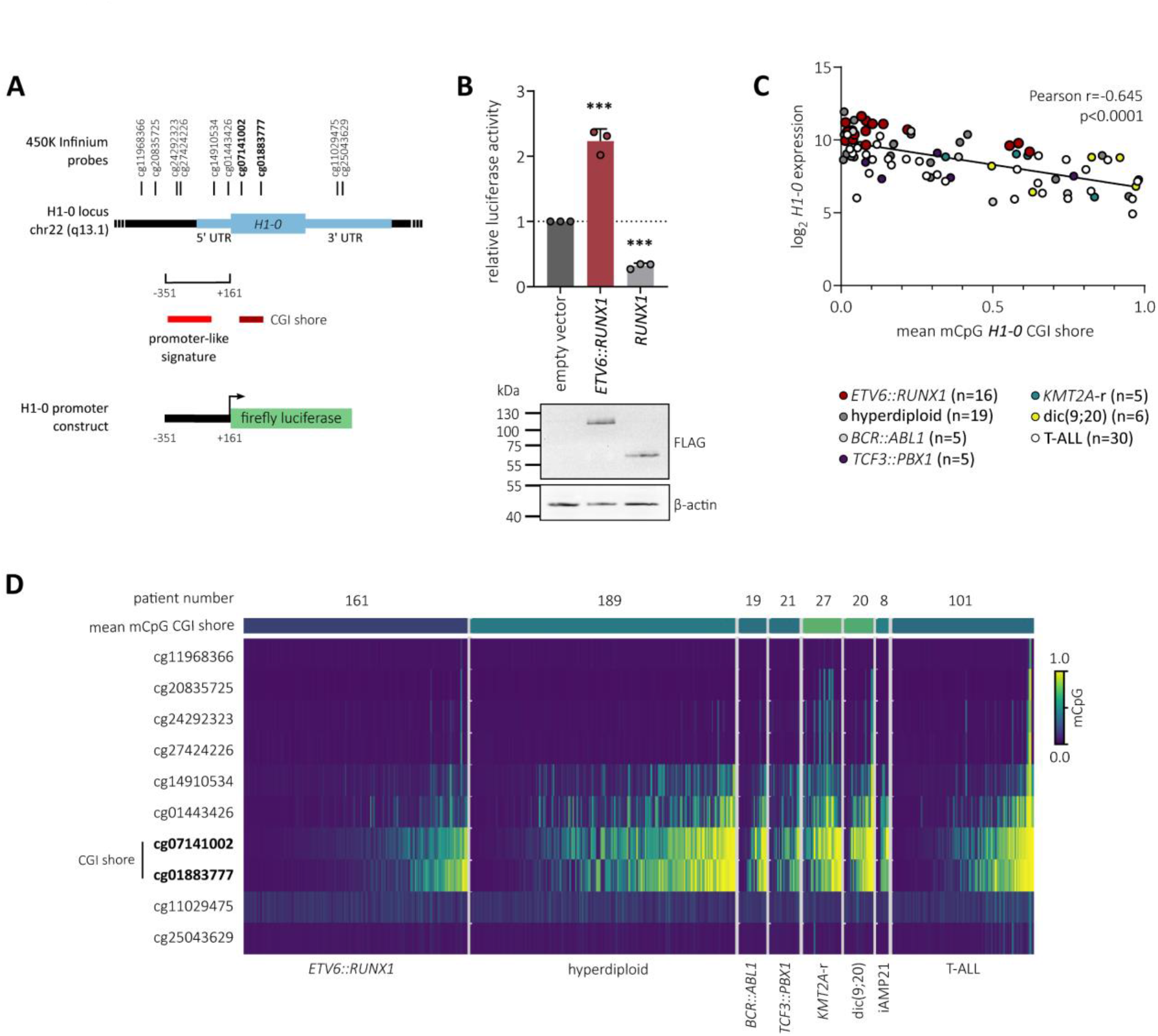
*ETV6::RUNX1* expression is sufficient to induce *H1-0* promoter activation. **(A)** Schematic representation of the *H1-0* locus, including the 512-bp region (nucleotides −351 to +161 from TSS) encompassing promoter-like signature EH38E2163184 (ENCODE). The *H1-0* CpG island (CGI) shore and 450K Infinium array probes are indicated. **(B)** 293T cells were transfected with a vector encoding the *H1-0* promoter-like signature indicated in (A), together with the empty pcDNA3.1 vector or pcDNA3.1 plasmids expressing either *ETV6::RUNX1* or *RUNX1*, and a vector expressing Renilla luciferase. Luciferase activities were normalized to Renilla luciferase activity and the empty vector control. Data represent mean values of three independent replicates ± standard deviation. Significance was calculated using an ordinary one-way ANOVA (***p<0.001). Representative protein levels of ETV6::RUNX1, RUNX1 and β-actin determined by Western blot are shown. **(C)** Pearson correlation of *H1-0* RNA expression and mean DNA methylation of the *H1-0* CGI shore probes cg07141002 and cg01883777 in leukemia patients (accession number GSE49032 (Nordlund et al, 2013)). Expression is shown for microarray probe 208886_at. Each dot represents a single patient. **(D)** *H1-0* DNA methylation in different leukemia entities is visualized as a heatmap with each row corresponding to a single patient (accession number GSE49032 (Nordlund et al, 2013)). Within each entity, patients are sorted according to mean DNA methylation of CGI shore probes cg07141002 and cg01883777. Total number of patients per entity is indicated.

Additionally to transcriptional control via binding of transcription factors, differential DNA methylation of the *H1-0* CpG island (CGI) shore has been reported to regulate *H1-0* expression in various solid tumor types, acting as an enhancer element (Morales Torres et al, 2016). Hence, we analyzed 450K Infinium microarray DNA methylation data comprising patient samples of T-ALL and six B-ALL subtypes (n=546, accession number GSE49032 (Nordlund et al, 2013)). Indeed, the average CGI shore methylation of *H1-0*, comprising probes cg07141002 and cg01883777, was lowest in *ETV6::RUNX1*+ BCP-ALL (Fig. 3C) and inversely correlated with *H1- 0* mRNA expression (Pearson r=-0.645, p<0.0001; Fig. 3D), indicating that *H1-0* expression is regulated via dynamic methylation of its CGI shore in leukemia.

### *H1-0* expression anti-correlates with cellular transcriptional activity during hematopoiesis

During hematopoiesis, *H1-0* is expressed in undifferentiated, quiescent progenitor cells (Valiron & Gorka, 1997). To characterize *H1-0* expression during B lymphopoiesis, we analyzed published RNA-seq (accession number GSE115656 (Black et al, 2018)), expression microarray (accession number GSE24759 (Novershtern et al, 2011)) and scRNA-seq (accession number E-MTAB-7407 (Popescu et al, 2019)) datasets. Across these datasets, we observed a continuous decrease of *H1-0* expression during B cell development (Fig. 4A-C) and significant upregulation of *H1-0* in *ETV6::RUNX1*+ ALL cells (n=6) compared to HSCs and later B cell progenitor stages (Fig. 4A).

**Figure 4.**
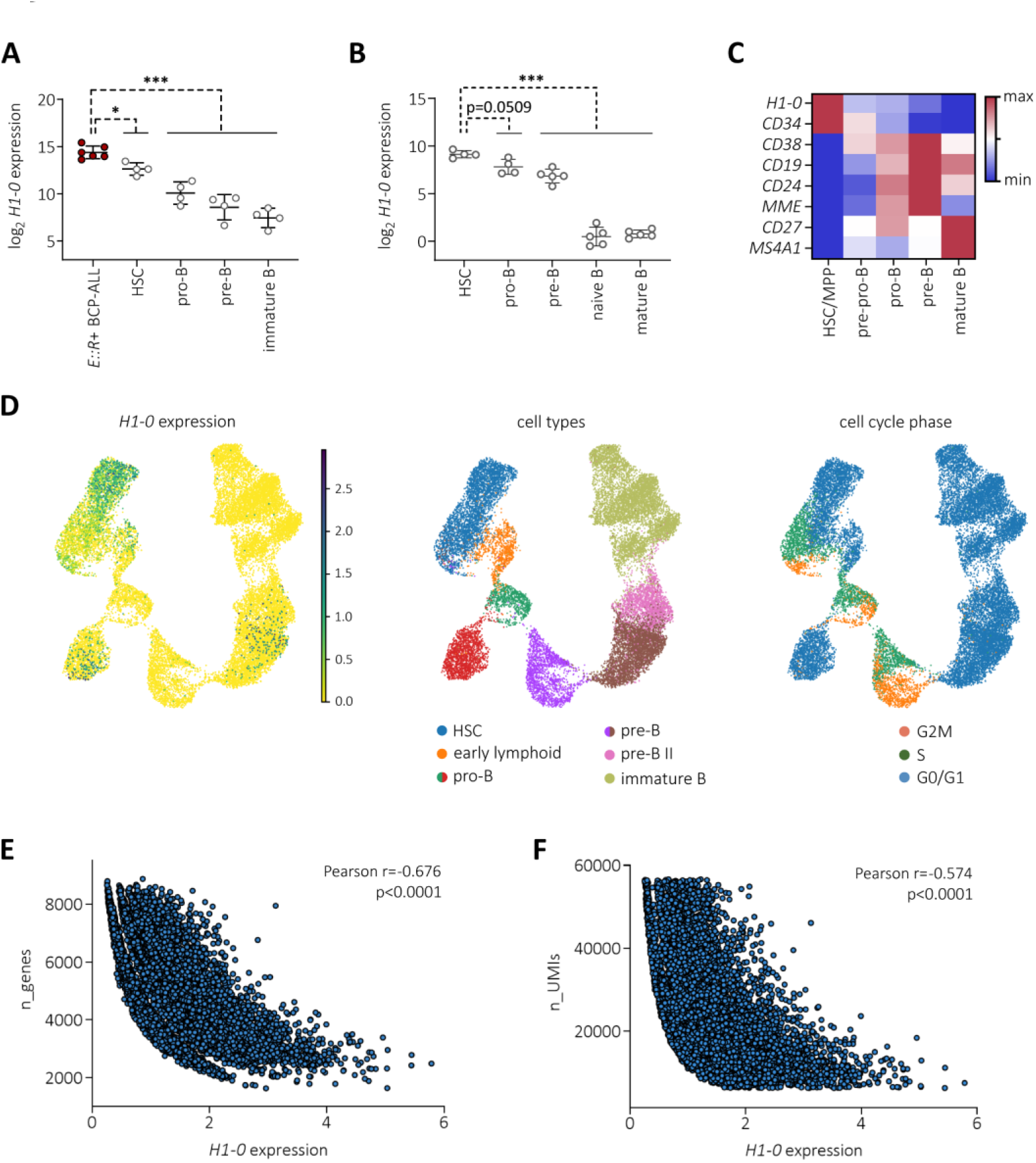
*H1-0* expression anti-correlates with cellular transcriptional activity during hematopoiesis. **(A)** *H1-0* expression in *ETV6::RUNX1*+ BCP-ALL (n=6) and healthy B cell precursor stages derived from a published RNA-seq dataset (accession number GSE115656 (Black et al, 2018)). B cell precursor fractions are HSCs (CD34+CD19-IgM-), pro-B cells (CD34+CD19+IgM-), pre-B cells (CD34-CD19+IgM-) and immature B cells (CD34-CD19+IgM+). **(B)** *H1-0* expression in healthy B cell precursor stages derived from a published expression microarray dataset (accession number GSE24759 (Novershtern et al, 2011)). B cell precursor fractions are HSCs (CD34+CD38-), pro-B cells (CD34+CD10+CD19+), pre-B cells (CD34- CD10+CD19+), naïve B cells (CD19+IgD+CD27-) and mature B cells (CD19+IgD+CD27+). (B, C) Mean expression ± standard deviation is indicated and data was analyzed for statistical significance using an ordinary one-way ANOVA (*p<0.05, ***p<0.001). **(C)** Min-max-normalized mean expression per cell type derived from a fetal liver scRNA-seq dataset (accession number E-MTAB-7407 (Popescu et al, 2019)). **(D)** *H1-0* expression levels across normal B-lymphoid differentiation distinguishing cell cycle status is depicted in a scRNA-seq UMAP visualization of B cell precursor cells from bone marrow of 8 healthy donors (Mehtonen et al, 2020). **(E, F)** Pearson correlation of *H1-0* expression (log-normalized counts) and total number of expressed genes (n_genes) or unique molecular identifier counts (n_UMIs) per cell detected by scRNA- seq in hiPSC-derived HPCs. *H1-0* counts of 0 were excluded.

To examine the expression level in context of cell cycle activity, we employed scRNA-seq data of B cell precursor cells derived from bone marrow of eight healthy donors and corresponding cell cycle and differentiation stage annotation (Mehtonen et al, 2020). *H1-0*+ cell numbers decreased along the B lineage trajectory, clustering preferentially to undifferentiated and G0/G1 cell cycle phases (Fig. 4D). Moreover, we found significant anti-correlation of *H1-0* expression with number of transcripts per cell (n_UMIs) and total number of genes per cell (n_genes, Fig. 4E-F). Indeed, transcriptional diversity (i.e. number of expressed genes per cell) is associated with cellular differentiation state (Gulati et al, 2020). Our data suggest that *H1-0* is an indicator of overall transcriptional activity and differentiation state.

### H1-0 is a key mediator of the ETV6::RUNX1-specific gene signature

To determine the contribution of *H1-0* to *ETV6::RUNX1*+ BCP-ALL pathology, we knocked down *H1-0* in the *ETV6::RUNX1*+ BCP-ALL cell line REH and performed gene set enrichment analysis (GSEA) on RNA-seq data. Knockdown resulted in ≈2.4-fold reduction of *H1-0* mRNA levels compared to non-targeting siRNA (siCtrl) treatment (Fig. 5A). GSEA using the canonical pathways collection (Human MSigDB Collections) revealed significant enrichment (cut-offs: p<0.005, false discovery rate (FDR) q-value<0.1) of gene signatures associated with DNA replication, histone modification, DNA repair and protein ubiquitination in siCtrl-treated REH cells (Fig. 5B and Dataset EV4), while no gene sets were identified as significantly enriched in REH cells treated with *H1-0*-targeting siRNA using the same cut-offs (Dataset EV5). Notably, GSEA detected enrichment of gene sets linked to histone acetylation (Dataset EV4 [in red]) in siCtrl-treated REH cells, consistent with previous reports highlighting strong correlation between *H1-0* gene expression and chromatin acetylation (Girardot et al, 1994; Morales Torres et al, 2020).

**Figure 5.**
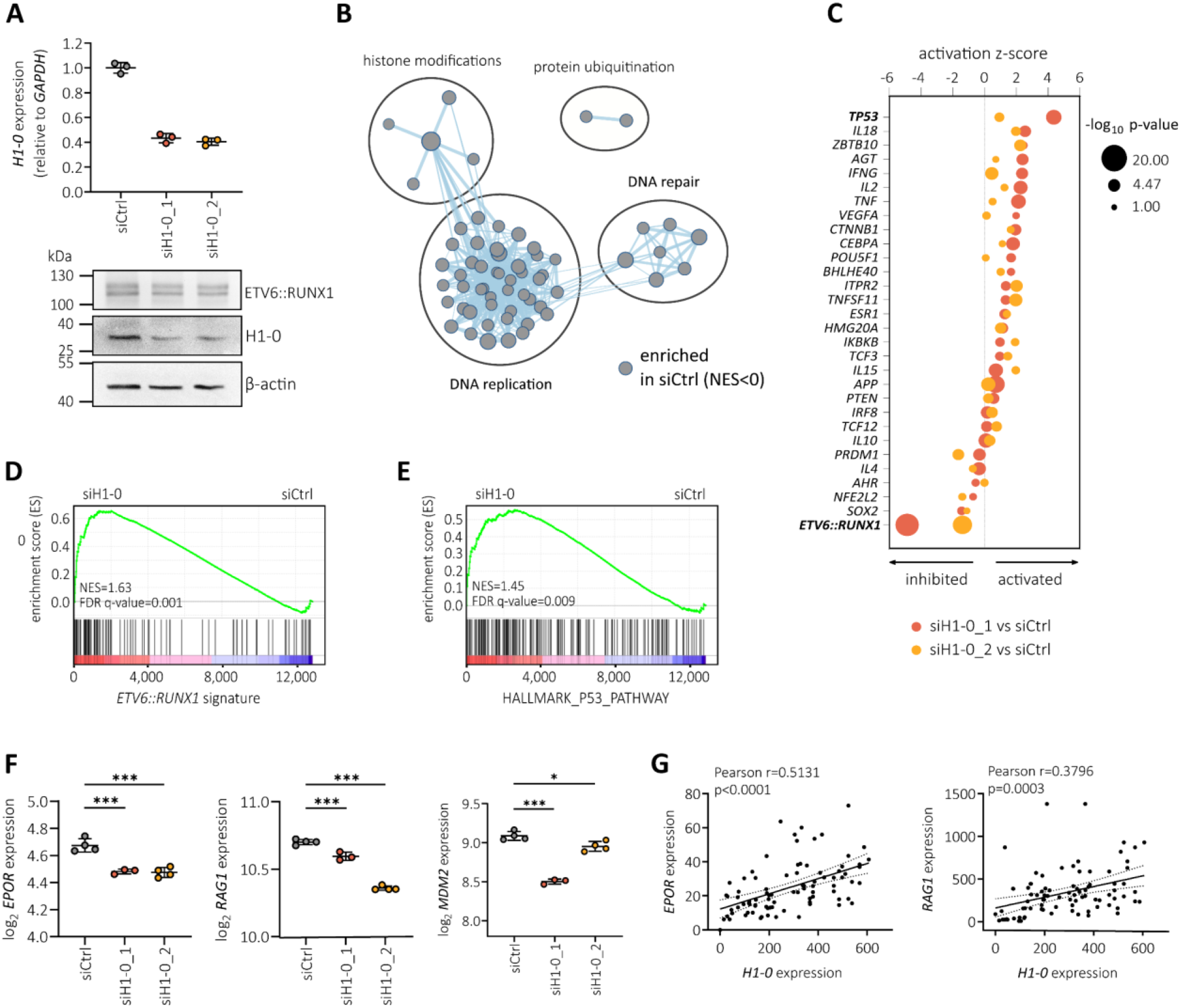
H1-0 is a key mediator of the ETV6::RUNX1-specific gene signature. **(A)** *H1-0* expression determined by RT-qPCR and representative Western Blot of REH cells treated for 48 hours with a non-targeting siRNA pool (siCtrl) or *H1-0*-targeting siRNA pools siH1-0_1 or siH1- 0_2. Data is presented as the mean ± standard deviation. **(B)** Enrichment map of gene sets enriched in siCtrl REH cells compared to siRNA-mediated knockdown of *H1-0* (cut-offs: p<0.005, false discovery rate (FDR) q-value<0.1) using the canonical pathways gene set collection (Human MSigDB Collections). No significantly enriched gene sets were found in siH1-0 REH cells using the indicated cut-offs. Groups of similar pathways are indicated. **(C)** Ingenuity Pathway analysis (IPA, QIAGEN) of upstream regulators significantly enriched in both siH1-0_1 versus siCtrl and siH1-0_2 versus siCtrl (p<0.05). **(D)** GSEA results of siH1-0 versus siCtrl using a published geneset (Fuka et al, 2011b) of 103 significantly upregulated genes in both REH and AT-2 cells upon *ETV6::RUNX1* knockdown (cut-offs: log2 fold change >0.9 and adjusted p<0.05). Normalized enrichment score (NES) and FDR are indicated. **(E)** GSEA of siH1-0 versus siCtrl using the HALLMARK_P53_PATHWAY gene set derived from Human MSigDB Collections. **(F)** RNA expression levels of *EPOR*, *RAG1* and *MDM2* determined by RNA-seq in siCtrl and siH1-0 REH. **(G)** Pearson correlation of *H1-0* expression with *EPOR* or *RAG1* expression in *ETV6::RUNX1*+ BCP-ALL patient samples derived from the PeCan St. Jude cloud (n=87, (Downing et al, 2012; McLeod et al, 2021)).

To ascertain common molecular drivers of gene expression changes observed upon *H1-0* knockdown, we applied upstream regulator analysis using the Ingenuity Pathway Analysis (IPA) suite (QIAGEN). Interestingly, the most significant potential driver detected by IPA upstream regulator analysis is ETV6::RUNX1, with p=3.7 x 10^-16^ (for siH1-0_1 vs. siCtrl, Dataset EV6) and p=3.5 x 10^-11^ (for siH1-0_2 vs. siCtrl, Dataset EV7) respectively (Fig. 5C). Negative activation z-scores indicate predicted inhibition of the ETV6::RUNX1 transcription factor upon *H1-0* knockdown. Given that ETV6::RUNX1 primarily functions as a repressor of RUNX1 regulated genes (Hiebert et al, 1996a), we employed a set of ETV6::RUNX1-regulated genes (cut-offs: log2 fold change >0.9 and p<0.05) (Fuka et al, 2011b) to validate our findings. Indeed, GSEA revealed significant upregulation of *ETV6::RUNX1* signature genes upon *H1-0* knockdown (normalized enrichment score [NES]=1.63, false-discovery rate [FDR] q-value=0.001; Fig. 5D).

Furthermore, we detected significant activation of *TP53* (encoding for p53) following *H1-0* knockdown, as indicated by both upstream regulator analysis (Fig. 5C) and GSEA (NES=1.45, FDR q-value=0.009; Fig. 5E). Indeed, previous studies have demonstrated that ETV6::RUNX1 suppresses p53 activity by upregulating *MDM2* (Kaindl et al, 2014), which we found to be downregulated in REH cells upon *H1-0* knockdown (Fig. 5F). Moreover, both *EPOR* and *RAG1*, two genes upregulated by ETV6::RUNX1 and key factors in *ETV6::RUNX1*+ BCP-ALL pathophysiology (Chen et al, 2021; Jakobczyk et al, 2022; Papaemmanuil et al, 2014; Torrano et al, 2011), exhibited reduced levels upon *H1-0* knockdown (Fig. 5F) as well as significant correlation with *H1-0* mRNA expression in *ETV6::RUNX1*+ BCP-ALL patient samples derived from the PeCan St. Jude cohort (n=87, Fig. 5G) (Downing et al, 2012; McLeod et al, 2021). Taken together, these data imply that linker histone *H1-0* is a key regulator of ETV6::RUNX1-induced expression changes.

### H1-0 inducer Quisinostat synergizes with frontline drugs in *ETV6::RUNX1*+ leukemic cells

Due to their cytostatic activity, HDACis are potent inducers of *H1-0* expression (Girardot et al, 1994; Morales Torres et al, 2020). Indeed, we found a striking correlation between H1-0 protein levels and drug sensitivity with HDACis in a panel of 25 BCP-ALL cell lines (Fig. 6A; data derived from the Functional Omics Resource of Acute Lymphoblastic Leukemia [FORALL] platform, https://proteomics.se/forall/ (Aswad & Jafari, 2023; Leo et al, 2022)), in particular with AR-42 and suberanilohydroxamic acid (SAHA)/Vorinostat (p<0.001), as well as 11 other HDACis, including Quisinostat (p<0.01). The pan-HDACi Quisinostat has been previously found to effectively upregulate *H1-0* expression and elicit arrest of cell proliferation at nanomolar concentrations in various solid cancer types (Morales Torres et al, 2020).

**Figure 6.**
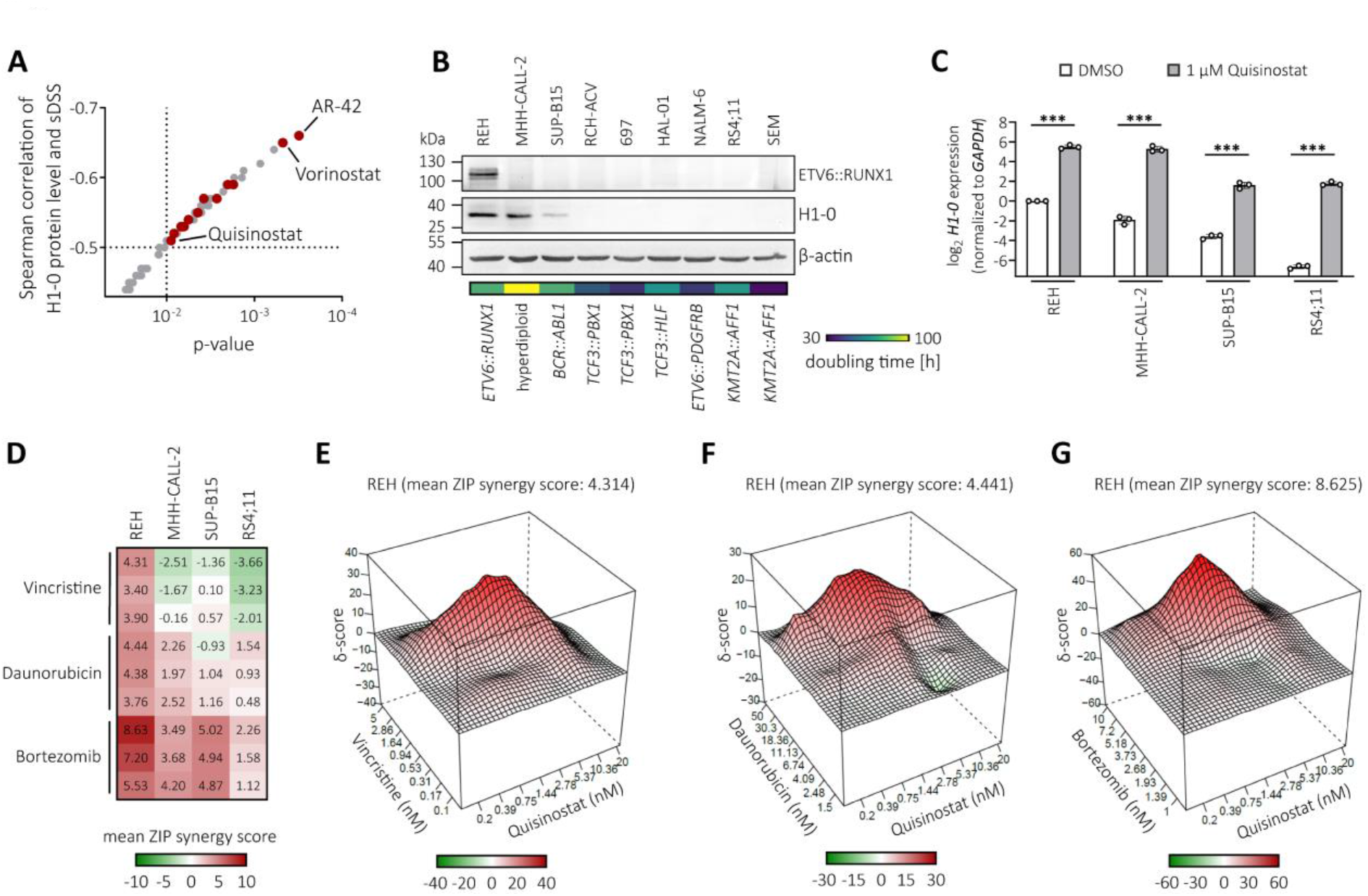
H1-0 inducer Quisinostat synergizes with frontline drugs in *ETV6::RUNX1*+ BCP-ALL. **(A)** Spearman correlation and Spearman p-values of H1-0 protein levels and selective drug sensitivity scores (sDSS) in 25 BCP-ALL cell lines derived from the FORALL platform (Aswad & Jafari, 2023; Leo et al, 2022) (https://proteomics.se/forall/; cut-offs: p<0.05, FDR<0.25). HDACis are marked in red. **(B)** Protein levels of ETV6::RUNX1, H1-0 and β-actin in BCP-ALL cell lines were quantified by Western blot. Chromosomal aberrations present in the respective cell line are indicated. Doubling times are derived from the DSMZ (https://www.dsmz.de/). **(C)** RT-qPCR quantifying *H1-0* levels 24 hours after treatment with DMSO or 1 µM Quisinostat in BCP-ALL cell lines. Values represent mean ± standard deviation from three independent replicates and data was analyzed for statistical significance using an ordinary one-way ANOVA (***p<0.001). **(D)** Heatmap indicating mean ZIP synergy scores of Vincristine (0.1-5 nM), Daunorubicin (1.5- 50 nM) or Bortezomib (1-10 nM) in combination with Quisinostat (0.2-20 nM) in four BCP-ALL cell lines treated for 72 hours. Data of three independent experiments is shown. Colors indicate synergy (red), lack of synergy (white) or antagonism (green). **(E-G)** Representative synergy plots visualizing drug combinations with high synergy in REH cells.

Therefore, we assessed whether the addition of Quisinostat can potentiate cell death induced by Vincristine or Daunorubicin, two commonly used chemotherapeutics in B-ALL treatment protocols (Pui et al, 2011). Western blot analysis of H1-0 protein levels in BCP-ALL cell lines indicated high expression in the *ETV6::RUNX1*+ cell line REH as well as the hyperdiploid cell line MHH-CALL-2 (derived from a near-haploid clone), while cell lines bearing other chromosomal translocations displayed low (SUP-B15) or undetectable levels (Fig. 6B). As anticipated, H1-0 levels reflected doubling times (Fig. 6B). Upon treatment with 1 µM Quisinostat for 24 hours, BCP-ALL cell lines with high (REH and MHH-CALL-2), medium (SUP-B15) and low *H1-0* expression (RS4;11) strongly upregulated *H1-0* (Fig. 6C). Combination of Quisinostat with the microtubule-depolymerizing agent Vincristine or the topoisomerase II inhibitor Daunorubicin, showed synergism in REH cells (Fig. 6D-F), whereas either no additive effect or a less pronounced additive effect was observed in MHH-CALL-2, SUP-B15 and RS4;11 cells (Fig. 6D). Simultaneous inhibition of HDACs and the proteasome has shown clinical efficacy in previous studies targeting hematologic malignancies (Bhatia et al, 2018). We detected high synergistic activity of Quisinostat with the proteasome inhibitor Bortezomib in all four tested BCP-ALL cell lines (Fig. 6D), particularly in REH cells (ZIP synergy score >30, Fig. 6G).

## Discussion

In this study, we established two independent preleukemic *ETV6::RUNX1*+ knock-in hiPSC models. Transcriptome analysis of these models revealed that ETV6::RUNX1 expression upregulates H1-0, a variant of the H1 linker histone family that promotes chromatin compaction (Hu et al, 2024; Willcockson et al, 2021). We demonstrate that H1-0 expression is associated with transcriptional activity, and significantly contributes to the repressive gene expression signature conferred by ETV6::RUNX1 as well as to p53 pathway activation. Attenuation of p53 has previously been connected to ETV6::RUNX1-mediated upregulation of MDM2 (Kaindl et al, 2014), which we found to be downregulated upon *H1-0* depletion.

High H1-0 levels observed in HSCs are in line with the largely quiescent nature of these cells (Valiron & Gorka, 1997). The progressive decrease of *H1-0* levels during hematopoiesis supports the notion that H1-0 accumulates in quiescent cells that have high proliferative capacity (Ebinger et al, 2016; Morales Torres et al, 2016). Increased quiescence of *ETV6::RUNX1*+ preleukemic cells is in keeping with the presence of these cells in the cord blood of approximately 5% of healthy newborns that we detected previously (Schäfer et al, 2018) and might offer an explanation for prolonged latency periods of *ETV6::RUNX1*+ leukemia of up to 14 years (Wiemels et al, 1999).

Interestingly, increased H1-0 levels were also observed in leukemic BCP-ALL patient samples, suggesting retention of chromatin compaction throughout *ETV6::RUNX1*+ BCP-ALL development. This is consistent with a recent report of reduced global chromatin accessibility in *ETV6::RUNX1*+ BCP-ALL compared to other ALL subtypes (Barnett et al, 2023). Similar loss of chromatin accessibility and cell cycle arrest has been detected in myeloid progenitors harboring the *RUNX1::ETO* translocation that retains the DNA-binding RHD, allowing it to bind to RUNX1 target sites (Nafria et al, 2020).

Clinically, the H1-0 inducer Quisinostat has demonstrated high potency and bioavailability at low nanomolar concentrations (Arts et al, 2009; Venugopal et al, 2013), while preserving normal stem cell function (Araki et al, 2006; Morales Torres et al, 2020; Young et al, 2004). However, the predominantly cytostatic activity of HDACis *in vivo* suggests that single-drug treatment is not sufficient to induce cancer remission. Using leukemic cell line models, we show here that combinatorial treatment using the pan-HDACi Quisinostat is a promising approach to enhance treatment of *ETV6::RUNX1*+ BCP-ALL when administered alongside Vincristine or Daunorubicin. Notably, we observed that combination of the proteasome inhibitor Bortezomib with Quisinostat demonstrated remarkable synergy in targeting *ETV6::RUNX1*+ leukemic cells at nanomolar concentrations. Indeed, combination of Bortezomib with Quisinostat showed promising treatment outcomes in a multiple myeloma mouse model (Deleu et al, 2009), while a previous study (Bastian et al, 2013) also reported efficacy of other pan-HDACis used in combination with Bortezomib in preclinical B-ALL models. As shown for solid cancer (Morales Torres et al, 2020), we propose that H1-0 levels indicate susceptibility towards HDACis in BCP-ALL. Therefore, combination treatments using Quisinostat might be a promising treatment approach for BCP-ALL, particularly for subtypes with high basal levels of the H1-0.

In conclusion, our data demonstrate that H1-0 contributes to the repressive function of ETV6::RUNX1 and corresponds to transcriptional activity in hematopoietic cells. Unraveling mechanisms involved in quiescence of *ETV6::RUNX1*+ preleukemic cells may offer new opportunities for enhancing patient treatment.

## Methods

### Cell lines and patient-derived xenografts

HW8 hiPSCs were generated from peripheral blood mononuclear cells (PBMCs) of a glioma patient using the CytoTune-iPS 2.0 Sendai Reprogramming kit (Thermo Fisher Scientific, #A16517) following written informed consent. Study approval was obtained by the internal review board at the National Institutes of Health (NIH, protocol number: 16CN 069). Cellartis human iPSC line 12 (ChiPSC12, #Y00280) was purchased from Takara Bio. BCP-ALL cell lines were obtained from the German Collection of Microorganisms and Cell Cultures (DSMZ). In-house leukemia patient samples for injection into NSG mice (The Jackson Laboratory) were retrieved from the Biobank of the University Hospital of Düsseldorf following informed consent in accordance with the Declaration of Helsinki. Study approval was obtained by the ethics committee of the Medical Faculty of the Heinrich Heine University (study number: 2019-566). All animal experiments adhered to regulatory guidelines set by the official committee at LANUV (Akt. 81-02.04.2017.A441) and were authorized by the animal research institute (ZETT) at Heinrich Heine University Düsseldorf. Patient blasts were injected intravenously into 6-week-old NSG mice (The Jackson Laboratory) and engraftment was assessed regularly by flow cytometric detection of human CD45+ cell percentage (BioLegend, #304011) in peripheral blood starting four weeks post injection (Vogt et al, 2023). Mice were sacrificed at predetermined timepoints and human CD45+ cells were isolated from bone marrow and spleen using the mouse cell depletion kit (Miltenyi Biotec, #130-104-694) to achieve >90% purity of human cells.

### Cell culture

hiPSCs were cultured at 37 °C and 5% CO2 on Geltrex-coated (Thermo Fisher Scientific, #A1413301) culture plates in mTeSR Plus (Stemcell Technologies, #100-0276) and were passaged in aggregates every 3-5 days using Versene solution (Thermo Fisher Scientific, #15040066) according to the manufacturer’s instructions. hiPSC lines were cultured for a maximum of 40 passages. Colony morphology was examined by microscopy (Axiovert 200, Zeiss). Pluripotency marker expression was assessed every 10 passages by flow cytometry (PE anti-human SSEA-4 antibody, BioLegend, #330405) or RT-qPCR for *DNMT3B*, *GDF3*, *POU5F1* and *NANOG*. Chromosomal integrity of hiPSCs was confirmed by karyotyping at the Institute of Human Genetics (Hannover Medical School (MHH), Germany).

BCP-ALL cell lines were maintained at 37 °C and 5% CO2 in RPMI GlutaMAX (Thermo Fisher Scientific, #61870036) with 20% fetal bovine serum (Biowest, #S181H-500). All cell lines were routinely tested for mycoplasma using the Venor GeM Advance mycoplasma detection kit (Minerva Biolabs, #11-7024). Short tandem repeat-based authentication of cell lines was performed at the Genomics and Transcriptomics Laboratory (GTL, Heinrich Heine University Düsseldorf, Germany) using the GenePrint 10 system (Promega, #B9510).

### Molecular cloning of a *RUNX1* HDR template

A *RUNX1* homology-directed repair (HDR) template targeting *ETV6* intron 5 was constructed by combining *RUNX1* exon sequences 2-8 with a puromycin resistance gene under control of the human *EF-1*α promoter. Homology arm sequences of ≈500 bp were PCR-amplified from ChiPSC12 genomic DNA and ligated to both sides of the HDR template. Single guide RNA (sgRNA) sequences targeting the 5’ region of *ETV6* intron 5 were designed using the online prediction tool CRISPOR (http://crispor.tefor.net/) (Concordet & Haeussler, 2018) and subcloned into the pUC19-U6-BbsI-sgRNA plasmid. Ligation of sequences was performed by PCR-based cloning via addition of specific restriction sites using the Phusion High-Fidelity PCR Master Mix (New England Biolabs, #F531L), restriction digest and subsequent ligation using T4 DNA ligase (New England Biolabs, #M0202). The *RUNX1* HDR template was subcloned into plasmid pUC19 (Addgene, #50005), linearized by PCR and concentrated by isopropanol precipitation to achieve a concentration ≥1 µg/µl. Primer sequences for molecular cloning were purchased from Eurofins Genomics and are listed in Dataset EV8. Target sequences selected for CRISPR/Cas9 genome editing of hiPSCs were GGATGAGGCTAAATCCCTAA (hg38, chr12: 11,870,115-11,870,134, + strand) and GCCTAATTGGGAATGGTGCG (hg38, chr12: 11870054-11870073, - strand).

### CRISPR/Cas9-editing of hiPSCs

Following incubation with 10 µM Y-27632 (STEMCELL Technologies, #72304) for 2 hours, single-cell suspensions of HW8 or ChiPSC12 hiPSCs were prepared using StemPro Accutase (Thermo Fisher Scientific, #A1110501). 10 x 10^6^ hiPSCs were resuspended in 100 µl P3 solution with supplement (Lonza, #V4XP-3024) and transfected with 2.5 µg linearized *RUNX1* HDR template and 4 µg each of pCW-Cas9 plasmid (Addgene, #50661) as well as two sgRNA plasmids using program CD-118 of the 4D Nucleofector system (Lonza). hiPSCs were plated onto 10 cm Geltrex-coated dishes in mTeSR Plus/Y-27632. Medium was exchanged to mTeSR Plus without Y-27632 after 24 h. Selection with 0.5 µg/ml puromycin (Thermo Fisher Scientific, #A1113803) was commenced 48 h after Nucleofection and single colonies were picked under microscopic guidance into a 96-well plate at day 7-10. Clones were expanded for subsequent confirmation of correct HDR template insertion.

Genomic DNA was isolated from puromycin-resistant hiPSC colonies using the QIAamp DNA Blood Mini kit (Qiagen, #51104). To ensure correct incorporation of the *RUNX1* HDR template into the *ETV6* locus, genotyping PCRs were performed with two sets of primers (5’-fw and 5’-rev, 3’-fw and 3’-rev). To confirm heterozygous allele status of *ETV6::RUNX1*, a third PCR combining primers 5’-fw, 3’-rev and 5’-fw2 was performed. Primer sequences are listed in Dataset EV8. Genotyping results were confirmed by Sanger sequencing, performed at the GTL (Heinrich Heine University Düsseldorf, Germany).

### *In vitro* differentiation of hiPSCs

To ensure reproducibility, feeder-free differentiation of hiPSCs was performed using the STEMdiff Hematopoietic kit (Stemcell Technologies, #05310) according to the manufacturer’s instructions. On day 12, suspension cells were harvested and used for downstream analyses.

### Quantitative real-time PCR (RT-qPCR)

RNA was extracted with the RNeasy Mini kit (Qiagen, #74104) according to the manufacturer’s instructions. On column DNA digestion was performed using the RNase free DNase Set (Qiagen, #79254). For RT-qPCR detection of *H1-0*, an additional DNase digest was performed with the TURBO DNA-free kit (Thermo Fisher Scientific, #AM1907). cDNA was synthesized from 1 µg of total RNA using M-MLV reverse transcriptase (Promega, #M1701) according to the manufacturer’s instructions. Due to limited amount of RNA following siRNA mediated *H1-0* knockdown in REH cells, 200 ng of total RNA were reverse transcribed. RT-qPCR was performed in triplicates using SYBR Green PCR Mix (Thermo Fisher Scientific, #4309155) or TaqMan Universal PCR Master Mix (Thermo Fisher Scientific, #4304437) and fluorescence was measured with the CFX384 Touch Real-Time PCR Detection System (Bio-Rad). *GAPDH*, *ATP5PB* and *PGK1* served as housekeeping genes for normalization and have previously been confirmed as top-ranking housekeeping genes during iPSC reprogramming (Panina et al, 2018). A list of RT-qPCR primers used in this study can be found in Dataset EV8. Relative quantification of expression was calculated using the ΔΔCt method. RT-qPCR primer efficiencies were validated by performing dilution series (85-110% efficiency, R2>0.98) and melt curve analyses (Taylor et al, 2019). No-template control (NTC) and no-reverse-transcriptase controls (NRT) were included on each plate.

### siRNA-mediated *H1-0* knockdown

Specific siRNA sequences for knockdown of *H1-0* in REH cells were designed using the Eurofins siRNA design tool (https://eurofinsgenomics.eu/en/ecom/tools/sirna-design/) and purchased from Eurofins Genomics. Sequences are listed in Dataset EV9. 1 x 10^6^ REH cells were transfected with 200 pmol of each siRNA pool (siCtrl, siH1-0_1, siH1-0_2) in the 100 µl nucleocuvette format using the 4D Nucleofector system (Lonza, SF solution, program DS-150).

### Bulk RNA sequencing and data analysis

HW8 and ChiPSC12 hiPSCs of similar passage number (±2 passages) were lysed on ice using RLT buffer (Qiagen, #79216) and detached by scraping. RNA was isolated using the RNeasy Mini kit (Qiagen, #74104) with on-column DNA digestion (Qiagen, #79254). REH cells were transfected with siRNA as described above and RNA was extracted after 48 hours. RNA quality was assessed with the 2100 Bioanalyzer system (Agilent, #G2939BA). Sequencing was performed at the next-generation sequencing core facility of the German Cancer Research Center (DKFZ).

Barcoded libraries were prepared from 0.5 µg of total RNA using the TruSeq RNA Sample Preparation v2 kit (low-throughput protocol, Illumina) and quantified with the Bioanalyzer system (Agilent). 7.5 pM denatured libraries were used as input for cBot (Illumina) and subjected to deep sequencing using the NovaSeq 6000 (Illumina) for 101 cycles, with an additional 7 cycles for index reading.

Analysis of fastq files was performed by using the Partek Flow software (Partek Incorporated, St. Louis, MO, USA). After assessing the read quality, a trimming step was performed (both ends: 13 bases at the 5’ end and 1 base at the 3’ end). One technical replicate of siH1-0_1 was excluded from the analysis since it did not pass quality control. After trimming, reads were aligned to the hg38 genome using the STAR v2.4.1d aligner. Unaligned reads were further processed using Bowtie 2 v2.2.5 aligner. Aligned reads were combined and expression was quantified against the Ensembl database (release 84) by the Partek Expectation-Maximization algorithm (Xing et al, 2006). Partek flow default settings were used in all analyses.

Unsupervised hierarchical clustering and heatmap visualization of samples was performed after normalizing mean expression to 0 with a standard deviation of 1 and using Pearson’s dissimilarity algorithm and average linkage in Partek Genomics Suite (Partek Incorporated). Ingenuity pathway analysis (IPA, Qiagen) of REH cells was performed taking into account significantly dysregulated genes (absolute fold change >1.5 and p<0.05). The significance cut-off for IPA to identify upstream regulators was set to p<0.05. Visualization of pathway networks was performed using the Cytoscape EnrichmentMap and AutoAnnotate applications (Reimand et al, 2019; Shannon et al, 2003).

Gene set enrichment analysis (GSEA) was performed on processed RNA-seq data of REH cells treated with non-targeting or *H1-0*-targeting siRNA pools using the GSEA v4.2.3 software (http://software.broadinstitute.org/gsea). Genes were ranked by the GSEA software using the signal-to-noise metric. The permutation type was set to gene_set and number of permutations to 5000. Canonical pathways or hallmark gene sets were obtained from the Molecular Signatures Database (MSigDB, https://www.gsea-msigdb.org). Genes significantly upregulated upon *ETV6::RUNX1* knockdown in REH and AT-2 cells (cutoffs: log2 fold change >0.9 and adjusted p<0.05, n=103 genes) were derived from a published dataset (Fuka et al, 2011a).

### Single-cell RNA sequencing and data analysis

Following *in vitro* differentiation of hiPSCs, hematopoietic progenitor cells of 5 wells were pooled, filtered and resuspended in PBS with 0.04% BSA (Biowest). Cell viability was determined to be >70% using the BD Rhapsody Single-Cell Analysis system (BD Biosciences). Single-cell suspensions were used for single-cell droplet library generation on the 10X Chromium Controller system using the Chromium Single Cell 3’ NextGEM Reagent kit v3.1 (10X Genomics) according to the manufacturer’s instructions. Sequencing was performed on a NextSeq 2000 system (Illumina). All scRNA-seq reactions were performed at the GTL (Heinrich Heine University Düsseldorf).

Raw sequencing data was processed using the 10X Genomics CellRanger software (v6.0.2). Raw BCL-files were demultiplexed and processed to fastq files using the CellRanger mkfastq pipeline. Alignment of reads and UMI counting was performed via the CellRanger count pipeline to generate a gene-barcode matrix (genome version: GRCh38, Ensemble release 98). The CellRanger aggr pipeline was used for aggregation and sequencing depth normalization.

Filtered cells were normalized using the PFlog1pPF method (Booeshaghi et al, 2022). Thus, differences in sequencing depth were normalized via the normalize_total function in Scanpy (version 1.9.1) followed by variance stabilizing log+1 transformation by applying the log1p function, and a second depth normalization. Feature selection based on binomial deviance (Townes et al, 2019) was performed on raw counts using the package Scry (version 1.10.0, https://doi.org/10.18129/B9.bioc.scry). Subsequently, the 4000 most deviant genes were used to compute principal components (PCs) via the pca function in Scanpy. The top 50 PCs were used to calculate the neighborhood graph via the neighbors function (n_neighbors=15), and Uniform Manifold Approximation and Projection (UMAP) was computed using the umap function.

Cell cycle phase was inferred by scoring the cell cycle gene set as defined by Tirosh *et al*. (Tirosh et al, 2016) (from https://github.com/scverse/scanpy_usage/tree/master/180209_cell_cycle) by applying the score_genes_cell_cycle function. For cell type annotation, gene sets underlying cell types defined by Jardine *et al*. (Jardine et al, 2021) were extracted via the rank_genes_groups_df function (pval_cutoff=0.05, log2fc_min=2) and used for scoring via the score_genes function. For visualization purposes, gene expression values were scaled using scale (max_value=10) function.

### Dual-luciferase reporter assay

Human *H1-0* promoter sequence (nucleotides −351 to +161 from TSS) was PCR-amplified from REH cell genomic DNA and inserted into Firefly luciferase vector pGL4.22 (Promega, #E6771) at KpnI and HindIII restriction sites. 293T cells at 50-70% confluency were transfected with 755 ng plasmid DNA using Xfect Transfection Reagent (Clontech Laboratories, #631317) in 24-well plates according to the manufacturer’s instructions. Each well was transfected with 500 ng pGL4.22 vector with or without *H1-0* promoter expression as well as 5 ng Renilla luciferase control plasmid pGL4.73 (Promega, #E6911), and 250 ng of the respective pcDNA3.1 vectors (Thermo Fisher Scientific, #V79020) for expression of *ETV6::RUNX1* or *RUNX1* or empty vector in triplicates. Cells were lysed after 48 hours with Passive Lysis buffer and luciferase signal was measured on a Tecan SPARK 10M reader using the Dual-Luciferase Reporter Assay System (Promega, #E1910). Firefly luciferase signal was normalized to Renilla luciferase activity. Adequate protein expression of ETV6::RUNX1 and RUNX1 was determined by Western blot.

### Western blot

Cell lysis was performed with RIPA lysis buffer (Thermo Fisher Scientific, #89900) supplemented with protease inhibitors (Roche, #11836145001). Protein concentration was determined using a bicinchoninic acid assay (BCA, Thermo Fisher Scientific, #23225), samples were mixed with 6X Laemmli buffer (Thermo Fisher Scientific, #J61337) and denatured at 95 °C for 5 min. 5-20 µg protein lysate were separated by SDS polyacrylamide gel electrophoresis (SDS-PAGE) using 8-10% gels and transferred to Amersham Protran 0.45 µm nitrocellulose membranes (Merck, #GE10600012) by wet blotting using the Mini-Protean Vertical Electrophoresis system (Bio-Rad). Membranes were blocked for 1 hour with 5% BSA (Merck, #A3294) in T-BST at room temperature and incubated with primary antibodies (ETV6::RUNX1: rabbit IgG monoclonal, 1:2000, Abcam, #ab92336; ETV6: mouse monoclonal IgG, 1:2000, Santa Cruz Biotechnology, #sc-166835; H1-0: rabbit monoclonal IgG, 1:2000, Thermo Fisher Scientific, #MA5-35484 (ARC1059); ACTB: mouse monoclonal IgG2a, 1:5000, Merck, #A5316; FLAG: mouse IgG1 monoclonal, M2, 1:2000, Sigma-Aldrich, #F1804) diluted in blocking buffer overnight at 4 °C. The next day, membranes were incubated with secondary antibodies (goat anti-rabbit monoclonal IgG, HRP-linked, 1:1000, Cell Signaling Technology, #7074S; horse anti-mouse IgG monoclonal, HRP-linked, 1:2000, Cell Signaling Technology, #7076S) diluted in blocking solution for 1 hour at room temperature. Signal development was performed using ECL detection reagent (Merck, #GERPN2109) according to the manufacturer’s instructions and images were acquired using the JESS Western system (Proteinsimple, Bio-Techne). For further detections, membranes were incubated in T-BST with added 0.1% NaN3 (Merck, #S2002) in T-BST for 1 h at room temperature or Re-Blot Plus Strong Antibody Stripping solution for 15 min at room temperature (Merck, #2504). Quantification of protein bands was performed using ImageJ analysis software (release 1.53c).

### Flow cytometry

Output of CD34+ cells following hematopoietic differentiation of hiPSCs was determined by staining with PE/Dazzle594-labeled anti-human CD34 antibodies (BioLegend, #343534). Prior to addition of antibodies, unspecific binding sites were blocked with human TruStain FcX blocking reagent (BioLegend, #422301) and dead cells were excluded by staining with Viobility 405/520 fixable dye (Miltenyi Biotec, #130109814).

### *In vitro* inhibitor treatments

BCP-ALL cell lines were treated with 1 µM JNJ-26481585/Quisinostat at a concentration of 1 x 10^6^ cells per ml and RNA was extracted after 24 hours for subsequent analysis of *H1-0* expression by RT-qPCR. DMSO-dissolved compounds were purchased from Selleck Chemicals and MedChem Express. For drug synergy analysis, Quisinostat (concentration range: 0.2 nM-20 nM), Vincristine (concentration range: 0.1 nM-5 nM), Daunorubicin (concentration range: 1.5 nM-50 nM) and Bortezomib (concentration range: 1 nM-10 nM) were printed in a randomized fashion in increasing concentrations in 8 x 8 matrices using a D300e digital dispenser (Tecan) and normalized with DMSO (Sigma-Aldrich, #2650). Plates were incubated for 72 hours and viability was determined by CellTiter-Glo Luminescent viability assay (Promega) using a Tecan SPARK 10M reader. Synergy scores were determined using the zero interaction potency method (ZIP) using the SynergyFinder web application (version 3.0).

### Bioinformatic analysis of data sets obtained from public resources

RNA-seq data of preleukemia models was obtained from the ArrayExpress functional genomics data collection website (https://www.ebi.ac.uk/biostudies/arrayexpress) accession number E-MTAB-6382(Böiers et al, 2018). RNA expression data of leukemia subtypes and normal B cell developmental stages was obtained from the St. Jude PeCan Data Portal (https://pecan.stjude.cloud) (Downing et al, 2012; McLeod et al, 2021) and from the R2 Genomics Analysis Visualization Platform (http://r2.amc.nl; GSE87070 dataset (Polak et al, 2019), microarray platform u133p2; GSE24759 dataset (Novershtern et al, 2011), microarray platform u133a). Processed DNA methylation (Infinium HumanMethylation450 BeadChip platform) and matched RNA expression data (microarray platform u133p2) of various leukemia entities was retrieved from the Gene Expression Omnibus database (NCBI GEO, https://www.ncbi.nlm.nih.gov/geo) accession number GSE49032 (Nordlund et al, 2013). Analysis of RNA expression data was performed as described for bulk RNA-seq. RNA-seq data of normal B cell developmental stages and *ETV6::RUNX1*+ BCP-ALL samples was retrieved from NCBI GEO (accession number GSE115656 (Black et al, 2018)) and processed using the Galaxy platform (https://usegalaxy.eu). Fastq files were trimmed using the Trimmomatic tool and aligned to the hg38 genome using the HISAT2 aligner. Expression was quantified using htseq-count against the UCSC database. To exclude effects of underlying predisposing syndromes, one *ETV6::RUNX1*+ BCP-ALL patient presenting with trisomy 21 was omitted from the analysis. Analysis of scRNA-seq data derived from normal bone marrow precursor B cells was performed as previously described (Mehtonen et al, 2020). Fetal liver scRNA-seq data was derived from the Developmental Cell Atlas accession number E-MTAB-7407 (Popescu et al, 2019) (Newcastle University, https://www.humancellatlas.org/). ChIP-seq datasets of REH cells for H3K4me1, H3K4me3, H3K27ac and RUNX1 (accession number GSE117684 (Jakobczyk et al, 2021)), as well as ETV6::RUNX1 (accession number GSE176084 (Jakobczyk et al, 2022)) were downloaded from NCBI GEO. Fastq files were processed as described for bulk RNA-seq and BAM files were visualized using IGV version 2.9.1 (https://igv.org) (Robinson et al, 2011).

## Statistical analysis

Statistical analysis of data was performed using GraphPad Prism version 9.5.1. The number (n) of replicates and statistical tests are indicated in the Fig. descriptions. Statistical significance was considered for p-values *p<0.05, **p<0.01 and ***p<0.001.

## Data availability

The datasets produced in this study are available in the following databases:

- RNA-seq data: Gene Expression Omnibus GSE270944
- scRNA-seq data: Gene Expression Omnibus GSE270945

## Authorship contributions

Vera H. Jepsen: Conceptualization; investigation; data curation; validation; methodology; formal analysis; visualization; writing – original draft; writing – review and editing. Andrea Hanel: Formal analysis; software; methodology; writing – review and editing. Daniel Picard: Formal analysis; software. Juha Mehtonen: Formal analysis. Rebecca Hasselmann: Investigation; formal analysis; visualization; writing – review and editing. Julian Schliehe-Diecks: Investigation; formal analysis; methodology. Katerina Scharov: Investigation. Jia-Wey Tu: Investigation. Rigveda Bhave: Resources. Ersen Kameri: Resources. Nan Qin: Resources. Herui Wang: Resources. Zhengping Zhuang: Resources. Rabea Wagener: Data curation; formal analysis. Lena Blümel: Resources. Tobias Lautwein: Formal analysis; software. Daniel Hein: Supervision; funding acquisition; conceptualization. Gesine Kögler: Resources. Marc Remke: Resources. Sanil Bhatia: Supervision; resources; writing – review and editing. Merja Heinäniemi: Supervision; writing – review and editing. Arndt Borkhardt: Funding acquisition; supervision; writing – review and editing. Ute Fischer: Funding acquisition; conceptualization; supervision; writing – review and editing.

## Acknowledgements

The authors thank Judith Bartel at the Institute of Human Genetics (Hannover Medical School (MHH), Germany) for performing karyotype analyses of hiPSC lines and David Koppstein for computational technical assistance. Computational infrastructure and support were provided by the Centre for Information and Media Technology (ZIM) at Heinrich Heine University Düsseldorf (Germany). This work was funded by German Research Foundation (495318549), Deutsche Krebshilfe (German Cancer Aid (70114736)), German José Carreras Leukemia Foundation (DJCLS 18R/2021) and the Parents’ initiative Löwenstern e.V..

## Disclosure and competing interest statement

The authors declare no competing interests.

---

**Figure EV1.**
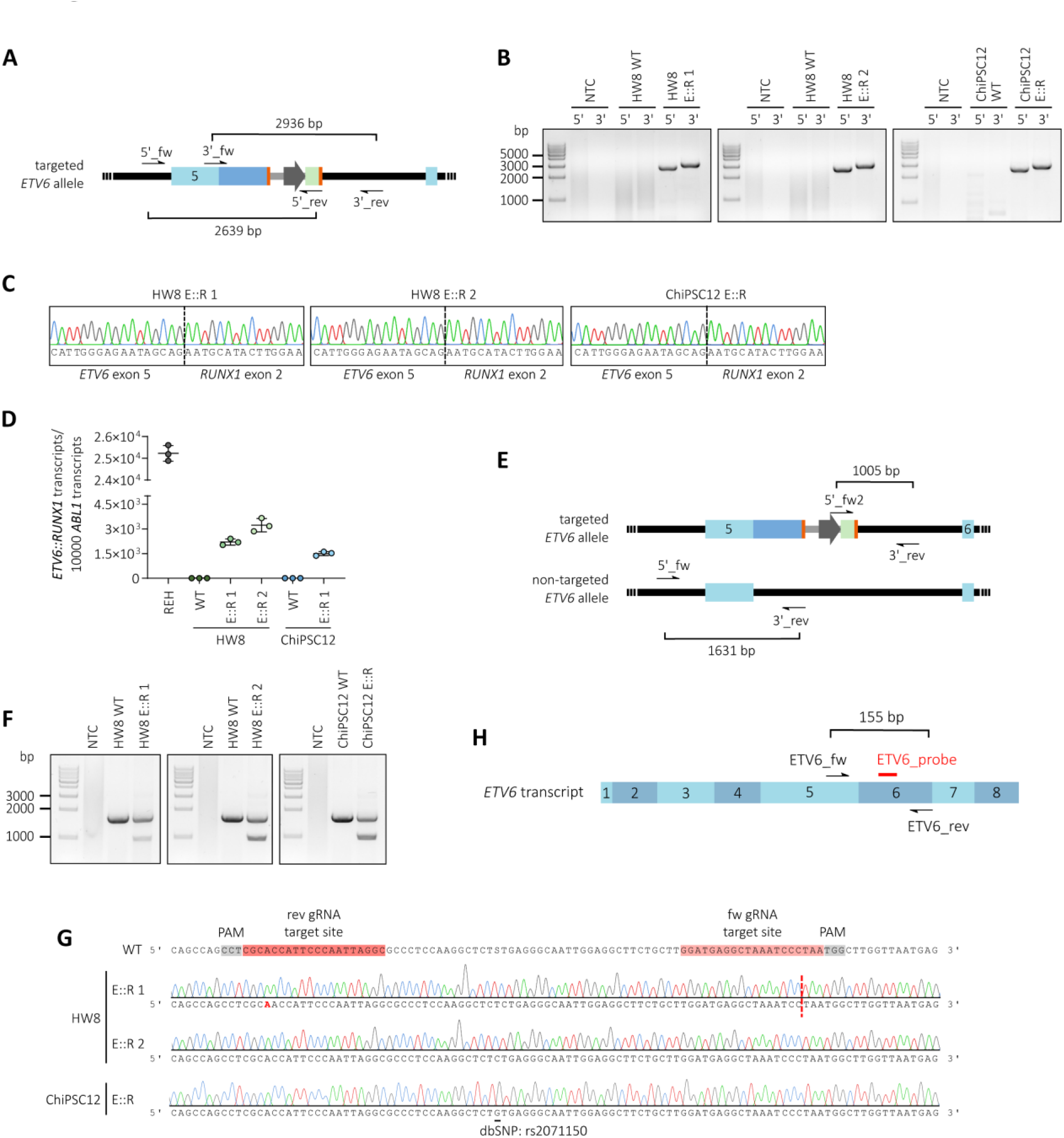
CRISPR/Cas9-edited HW8 and ChiPSC12 hiPSCs stably express *ETV6::RUNX1* from the endogenous *ETV6* locus. **(A)** Schematic representation of genotyping PCRs confirming correct insertion of the *RUNX1* HDR template into the *ETV6* locus of HW8 and ChiPSC12 hiPSCs (5’_PCR=2639 bp, 3’_PCR=2936 bp). Arrows indicate binding sites of PCR primers. **(B)** Genotyping PCRs confirming correct insertion of the *RUNX1* HDR template into the *ETV6* locus of HW8 and ChiPSC12 hiPSCs. **(C)** Sanger sequencing tracks of *ETV6::RUNX1*+ hiPSC lines detecting the fusion sequence between *ETV6* exon 5 and *RUNX1* exon 2. **(D)** Quantification of *ETV6::RUNX1* expression by RT-qPCR in the *ETV6::RUNX1*+ cell line REH, HW8 and ChiPSC12 hiPSC lines. Data is presented as the mean ± standard deviation. **(E)** Schematic representation of the PCR approach used for detection of *ETV6::RUNX1* homo- or heterozygosity in hiPSCs (WT allele=1631 bp, targeted allele=1005 bp). **(F)** PCRs confirming heterozygous allele status of *ETV6::RUNX1* in HW8 and ChiPSC12 hiPSCs. **(G)** Sanger sequencing of the gRNA binding region within *ETV6* intron 5 in the three *ETV6::RUNX1*+ hiPSC clones. **(H)** Schematic representation of the *ETV6* RT-qPCR design. The RT-qPCR probe sequence is marked in red. NTC: no-template control (nuclease-free H_2_O).

**Figure EV2.**
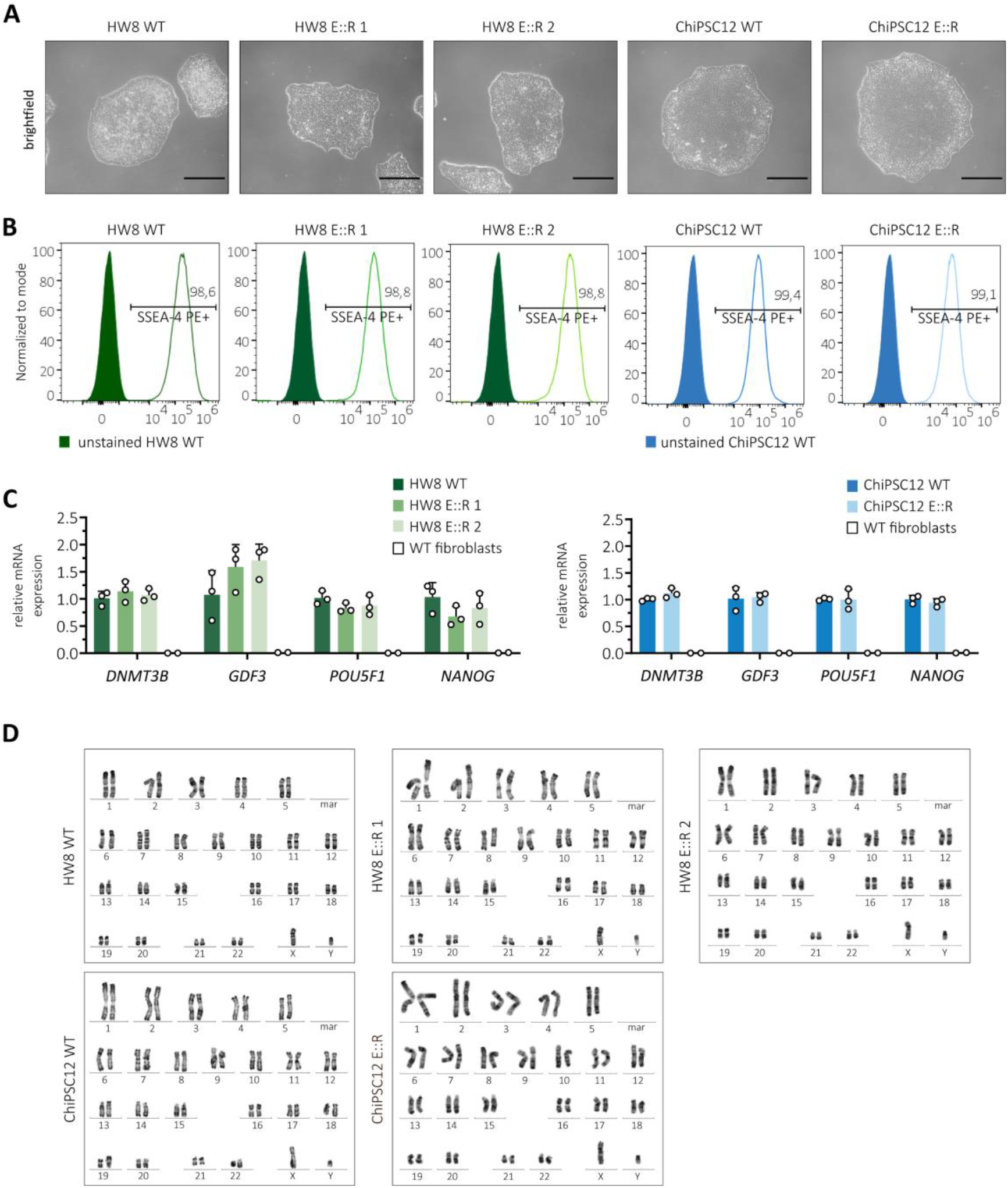
Assessment of hiPSC quality. **(A)** Representative brightfield images confirming normal hiPSC colony morphology (scale bar=300 µm). **(B)** Flow cytometric analysis of stage-specific embryonic antigen 4 (SSEA-4) on HW8 WT and ChiPSC12 WT cells, as well as CRISPR/Cas9-edited *ETV6::RUNX1*+ hiPSC lines HW8 E::R 1, HW8 E::R 2 and ChiPSC12 E::R. **(C)** Representative RT-qPCR analyses of pluripotency marker genes *DNMT3B*, *GDF3*, *POU5F1* and *NANOG* in HW8 WT, ChiPSC12 WT, as well as the respective ETV6::RUNX1+ hiPSC clones. Fibroblasts from two healthy donors were used as negative controls. Expression of *DNMT3B* and *GDF3* is normalized to *ATP5PB*, expression of *POU5F1* and *NANOG* is normalized to *PGK1*. Data is presented as the mean + standard deviation, and expression of *ETV6::RUNX1*+ hiPSC lines is presented relative to the respective WT. **(D)** Representative karyotype data for *ETV6::RUNX1*+ and WT hiPSCs. Karyotyping was performed by Judith Bartel at the Institute of Human Genetics (Hannover Medical School (MHH)).

**Figure EV3.**
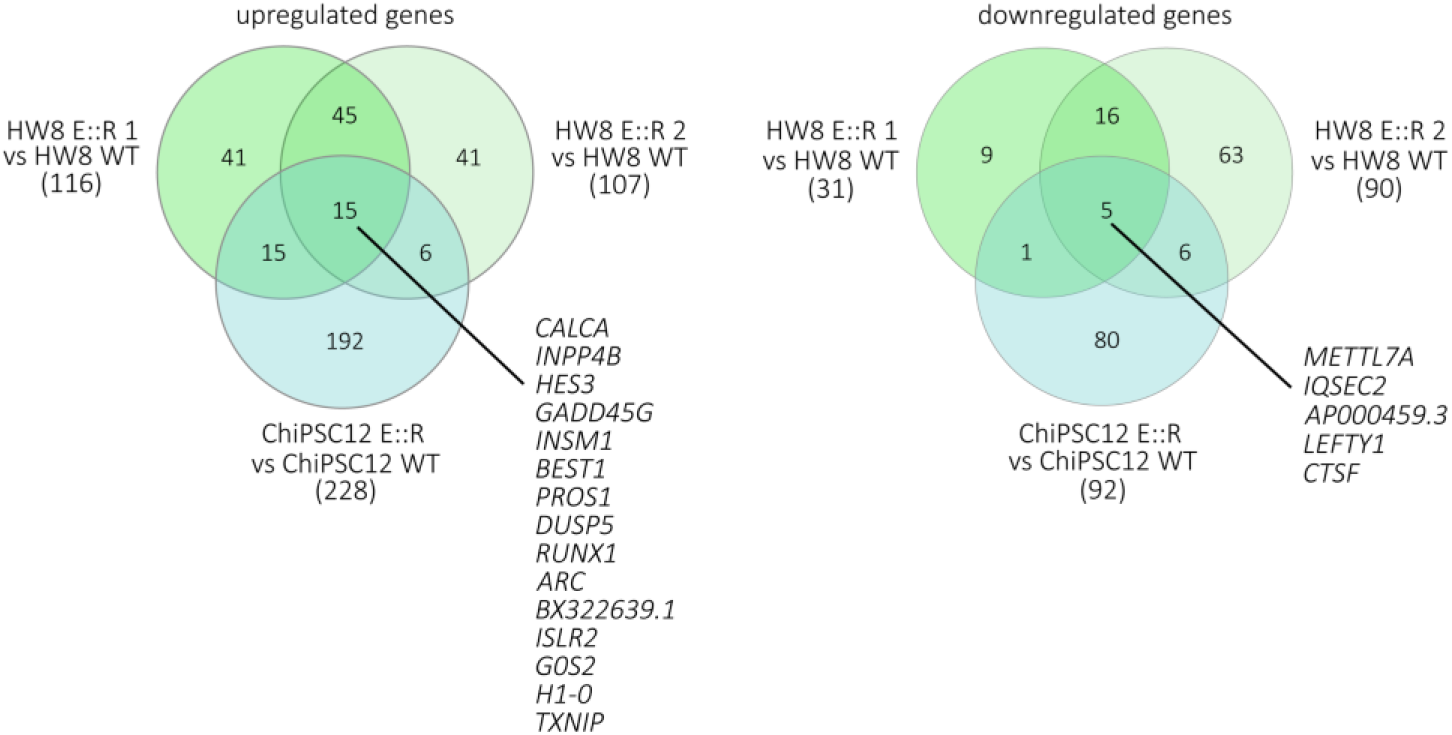
Transcriptome analysis of *ETV6::RUNX1*+ hiPSCs. Venn diagrams of upregulated and downregulated differentially expressed genes (absolute fold change >2 and p<0.05 detected in *ETV6::RUNX1*+ hiPSCs by RNA-seq compared to their respective WT counterpart.

**Figure EV4.**
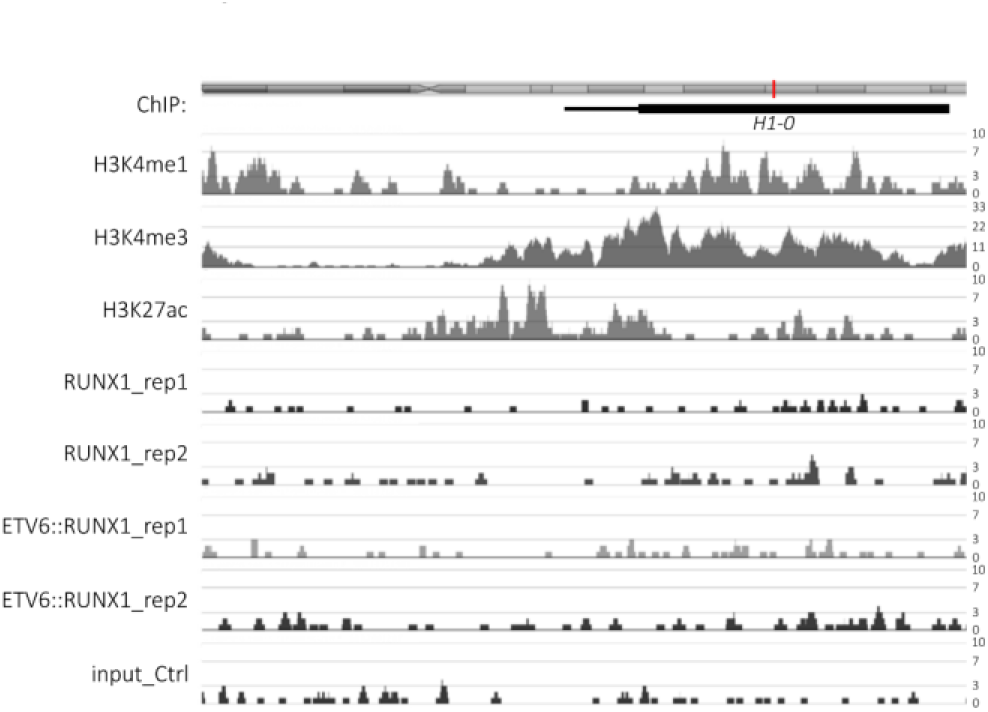
Chromatin immunoprecipitation (ChIP) analysis of the *H1-0* promoter region. ChIP peak visualization of the human *H1-0* gene region derived from ChIP-seq data of REH cells for H3K4me1, H3K4me3, H3K27ac and RUNX1 (accession number GSE117684 (Jakobczyk et al, 2021)) as well as ETV6::RUNX1 (accession number GSE176084 (Jakobczyk et al, 2022)).

## Notes

### Competing Interest Statement

The authors have declared no competing interest.

### Summary of Updates

Order of authors corrected; author affiliations updated; Supplementary Tables uploaded.

## References

1. Araki H, Yoshinaga K, Boccuni P, Zhao Y, Hoffman R, Mahmud N (2006) Chromatin-modifying agents permit human hematopoietic stem cells to undergo multiple cell divisions while retaining their repopulating potential. Blood 109: 3570–3578

2. Arts J, King P, Marin A, Floren W, Belin A, Janssen L, Pilatte I, Roux B, Decrane L, Gilissen R et al (2009) JNJ-26481585, a Novel Second-Generation Oral Histone Deacetylase Inhibitor, Shows Broad-Spectrum Preclinical Antitumoral Activity. Clinical Cancer Research 15: 6841–6851

3. Aswad L, Jafari R (2023) FORALL: an interactive shiny/R web portal to navigate multi-omics high-throughput data of pediatric acute lymphoblastic leukemia. Bioinformatics Advances 3

4. Barnett KR, Mobley RJ, Diedrich JD, Bergeron BP, Bhattarai KR, Monovich AC, Narina S, Yang W, Crews KR, Manring CS et al (2023) Epigenomic mapping reveals distinct B cell acute lymphoblastic leukemia chromatin architectures and regulators. Cell Genom 3: 100442

5. Bastian L, Hof J, Pfau M, Fichtner I, Eckert C, Henze G, Prada J, von Stackelberg A, Seeger K, Shalapour S (2013) Synergistic Activity of Bortezomib and HDACi in Preclinical Models of B-cell Precursor Acute Lymphoblastic Leukemia via Modulation of p53, PI3K/AKT, and NF-κB. Clinical Cancer Research 19: 1445–1457

6. Bernardin F, Yang Y, Cleaves R, Zahurak M, Cheng L, Civin CI, Friedman AD (2002) TEL-AML1, Expressed from t(12;21) in Human Acute Lymphocytic Leukemia, Induces Acute Leukemia in Mice1. Cancer Research 62: 3904–3908

7. Bhatia S, Krieger V, Groll M, Osko JD, Reßing N, Ahlert H, Borkhardt A, Kurz T, Christianson DW, Hauer J et al (2018) Discovery of the First-in-Class Dual Histone Deacetylase-Proteasome Inhibitor. J Med Chem 61: 10299–10309

8. Black KL, Naqvi AS, Asnani M, Hayer KE, Yang SY, Gillespie E, Bagashev A, Pillai V, Tasian SK, Gazzara MR et al (2018) Aberrant splicing in B-cell acute lymphoblastic leukemia. Nucleic Acids Res 46: 11357–11369

9. Böiers C, Richardson SE, Laycock E, Zriwil A, Turati VA, Brown J, Wray JP, Wang D, James C, Herrero J et al (2018) A Human IPS Model Implicates Embryonic B-Myeloid Fate Restriction as Developmental Susceptibility to B Acute Lymphoblastic Leukemia-Associated ETV6-RUNX1. Dev Cell 44: 362–377.e367

10. Booeshaghi AS, Hallgrímsdóttir IB, Gálvez-Merchán Á, Pachter L (2022) Depth normalization for single-cell genomics count data. bioRxiv: 2022.2005.2006.490859

11. Brady SW, Roberts KG, Gu Z, Shi L, Pounds S, Pei D, Cheng C, Dai Y, Devidas M, Qu C et al (2022) The genomic landscape of pediatric acute lymphoblastic leukemia. Nature Genetics 54: 1376–1389

12. Chen D, Camponeschi A, Nordlund J, Marincevic-Zuniga Y, Abrahamsson J, Lönnerholm G, Fogelstrand L, Mårtensson I-L (2021) RAG1 co-expression signature identifies ETV6-RUNX1- like B-cell precursor acute lymphoblastic leukemia in children. Cancer Medicine 10: 3997–4003

13. Concordet J-P, Haeussler M (2018) CRISPOR: intuitive guide selection for CRISPR/Cas9 genome editing experiments and screens. Nucleic Acids Research 46: W242–W245

14. Deleu S, Lemaire M, Arts J, Menu E, Van Valckenborgh E, Vande Broek I, De Raeve H, Coulton L, Van Camp B, Croucher P et al (2009) Bortezomib Alone or in Combination with the Histone Deacetylase Inhibitor JNJ-26481585: Effect on Myeloma Bone Disease in the 5T2MM Murine Model of Myeloma. Cancer Research 69: 5307–5311

15. Downing JR, Wilson RK, Zhang J, Mardis ER, Pui CH, Ding L, Ley TJ, Evans WE (2012) The Pediatric Cancer Genome Project. Nat Genet 44: 619–622

16. Ebinger S, Özdemir EZ, Ziegenhain C, Tiedt S, Castro Alves C, Grunert M, Dworzak M, Lutz C, Turati VA, Enver T et al (2016) Characterization of Rare, Dormant, and Therapy-Resistant Cells in Acute Lymphoblastic Leukemia. Cancer Cell 30: 849–862

17. Fenrick R, Amann JM, Lutterbach B, Wang L, Westendorf JJ, Downing JR, Hiebert SW (1999) Both TEL and AML-1 contribute repression domains to the t(12;21) fusion protein. Mol Cell Biol 19: 6566–6574

18. Ford AM, Palmi C, Bueno C, Hong D, Cardus P, Knight D, Cazzaniga G, Enver T, Greaves M (2009) The TEL-AML1 leukemia fusion gene dysregulates the TGF-beta pathway in early B lineage progenitor cells. J Clin Invest 119: 826–836

19. Fuka G, Kauer M, Kofler R, Haas OA, Panzer-Grumayer R (2011a) The leukemia-specific fusion gene ETV6/RUNX1 perturbs distinct key biological functions primarily by gene repression. PLoS One 6: e26348

20. Fuka G, Kauer M, Kofler R, Haas OA, Panzer-Grümayer R (2011b) The leukemia-specific fusion gene ETV6/RUNX1 perturbs distinct key biological functions primarily by gene repression. PloS one 6: e26348

21. Girardot V, Rabilloud T, Yoshida M, Beppu T, Lawrence J-J, Khochbin S (1994) Relationship between Core Histone Acetylation and Histone H10 Gene Activity. European Journal of Biochemistry 224: 885–892

22. Greaves M (2018) A causal mechanism for childhood acute lymphoblastic leukaemia. Nat Rev Cancer 18: 471–484

23. Gu Z, Churchman ML, Roberts KG, Moore I, Zhou X, Nakitandwe J, Hagiwara K, Pelletier S, Gingras S, Berns H et al (2019) PAX5-driven subtypes of B-progenitor acute lymphoblastic leukemia. Nat Genet 51: 296–307

24. Guidez F, Petrie K, Ford AM, Lu H, Bennett CA, MacGregor A, Hannemann J, Ito Y, Ghysdael J, Greaves M et al (2000) Recruitment of the nuclear receptor corepressor N-CoR by the TEL moiety of the childhood leukemia–associated TEL-AML1 oncoprotein. Blood 96: 2557–2561

25. Gulati GS, Sikandar SS, Wesche DJ, Manjunath A, Bharadwaj A, Berger MJ, Ilagan F, Kuo Angera H, Hsieh RW, Cai S et al (2020) Single-cell transcriptional diversity is a hallmark of developmental potential. Science 367: 405–411

26. Heumos L, Schaar AC, Lance C, Litinetskaya A, Drost F, Zappia L, Lücken MD, Strobl DC, Henao J, Curion F et al (2023) Best practices for single-cell analysis across modalities. Nature Reviews Genetics 24: 550–572

27. Hiebert SW, Sun W, Davis JN, Golub T, Shurtleff S, Buijs A, Downing JR, Grosveld G, Roussell M, Gilliland DG (1996a) The t (12; 21) translocation converts AML-1B from an activator to a repressor of transcription. Molecular and cellular biology

28. Hiebert SW, Sun W, Davis JN, Golub T, Shurtleff S, Buijs A, Downing JR, Grosveld G, Roussell MF, Gilliland DG et al (1996b) The t(12;21) translocation converts AML-1B from an activator to a repressor of transcription. Mol Cell Biol 16: 1349–1355

29. Hu S, Chapski DJ, Gehred ND, Kimball TH, Gromova T, Flores A, Rowat AC, Chen J, Packard RRS, Olszewski E et al (2024) Histone H1.0 couples cellular mechanical behaviors to chromatin structure. Nature Cardiovascular Research 3: 441–459

30. Jakobczyk H, Debaize L, Soubise B, Avner S, Rouger-Gaudichon J, Commet S, Jiang Y, Serandour AA, Rio AG, Carroll JS et al (2021) Reduction of RUNX1 transcription factor activity by a CBFA2T3-mimicking peptide: application to B cell precursor acute lymphoblastic leukemia. J Hematol Oncol 14: 47

31. Jakobczyk H, Jiang Y, Debaize L, Soubise B, Avner S, Sérandour AA, Rouger-Gaudichon J, Rio A-G, Carroll JS, Raslova H et al (2022) ETV6-RUNX1 and RUNX1 directly regulate RAG1 expression: one more step in the understanding of childhood B-cell acute lymphoblastic leukemia leukemogenesis. Leukemia 36: 549–554

32. Jardine L, Webb S, Goh I, Quiroga Londoño M, Reynolds G, Mather M, Olabi B, Stephenson E, Botting RA, Horsfall D et al (2021) Blood and immune development in human fetal bone marrow and Down syndrome. Nature 598: 327–331

33. Kaindl U, Morak M, Portsmouth C, Mecklenbräuker A, Kauer M, Zeginigg M, Attarbaschi A, Haas OA, Panzer-Grümayer R (2014) Blocking ETV6/RUNX1-induced MDM2 overexpression by Nutlin-3 reactivates p53 signaling in childhood leukemia. Leukemia 28: 600–608

34. Kodgule R, Goldman JW, Monovich AC, Saari T, Aguilar AR, Hall CN, Rajesh N, Gupta J, Chu SA, Ye L et al (2023) ETV6 Deficiency Unlocks ERG-Dependent Microsatellite Enhancers to Drive Aberrant Gene Activation in B-Lymphoblastic Leukemia. Blood Cancer Discov 4: 34–53

35. Konrad M, Metzler M, Panzer S, Ostreicher I, Peham M, Repp R, Haas OA, Gadner H, Panzer- Grumayer ER (2003) Late relapses evolve from slow-responding subclones in t(12;21)- positive acute lymphoblastic leukemia: evidence for the persistence of a preleukemic clone. Blood 101: 3635–3640

36. Leo IR, Aswad L, Stahl M, Kunold E, Post F, Erkers T, Struyf N, Mermelekas G, Joshi RN, Gracia-Villacampa E et al (2022) Integrative multi-omics and drug response profiling of childhood acute lymphoblastic leukemia cell lines. Nat Commun 13: 1691

37. Linka Y, Ginzel S, Borkhardt A, Landgraf P (2014) Identification of TEL-AML1 (ETV6-RUNX1) associated DNA and its impact on mRNA and protein output using ChIP, mRNA expression arrays and SILAC. Genom Data 2: 85–88

38. Linka Y, Ginzel S, Krüger M, Novosel A, Gombert M, Kremmer E, Harbott J, Thiele R, Borkhardt A, Landgraf P (2013) The impact of TEL-AML1 (ETV6-RUNX1) expression in precursor B cells and implications for leukaemia using three different genome-wide screening methods. Blood Cancer J 3: e151

39. McLeod C, Gout AM, Zhou X, Thrasher A, Rahbarinia D, Brady SW, Macias M, Birch K, Finkelstein D, Sunny J et al (2021) St. Jude Cloud: A Pediatric Cancer Genomic Data-Sharing Ecosystem. Cancer Discovery 11: 1082–1099

40. Mehtonen J, Teppo S, Lahnalampi M, Kokko A, Kaukonen R, Oksa L, Bouvy-Liivrand M, Malyukova A, Makinen A, Laukkanen S et al (2020) Single cell characterization of B-lymphoid differentiation and leukemic cell states during chemotherapy in ETV6-RUNX1-positive pediatric leukemia identifies drug-targetable transcription factor activities. Genome Med 12: 99

41. Morales Torres C, Biran A, Burney MJ, Patel H, Henser-Brownhill T, Cohen AS, Li Y, Ben-Hamo R, Nye E, Spencer-Dene B et al (2016) The linker histone H1.0 generates epigenetic and functional intratumor heterogeneity. Science 353

42. Morales Torres C, Wu MY, Hobor S, Wainwright EN, Martin MJ, Patel H, Grey W, Grönroos E, Howell S, Carvalho J et al (2020) Selective inhibition of cancer cell self-renewal through a Quisinostat-histone H1.0 axis. Nature Communications 11: 1792

43. Mori H, Colman SM, Xiao Z, Ford AM, Healy LE, Donaldson C, Hows JM, Navarrete C, Greaves M (2002) Chromosome translocations and covert leukemic clones are generated during normal fetal development. Proc Natl Acad Sci U S A 99: 8242–8247

44. Morrow M, Samanta A, Kioussis D, Brady HJ, Williams O (2007) TEL-AML1 preleukemic activity requires the DNA binding domain of AML1 and the dimerization and corepressor binding domains of TEL. Oncogene 26: 4404–4414

45. Mullighan CG, Goorha S, Radtke I, Miller CB, Coustan-Smith E, Dalton JD, Girtman K, Mathew S, Ma J, Pounds SB et al (2007) Genome-wide analysis of genetic alterations in acute lymphoblastic leukaemia. Nature 446: 758–764

46. Nafria M, Keane P, Ng ES, Stanley EG, Elefanty AG, Bonifer C (2020) Expression of RUNX1-ETO Rapidly Alters the Chromatin Landscape and Growth of Early Human Myeloid Precursor Cells. Cell Reports 31

47. Nordlund J, Bäcklin CL, Wahlberg P, Busche S, Berglund EC, Eloranta M-L, Flaegstad T, Forestier E, Frost B-M, Harila-Saari A et al (2013) Genome-wide signatures of differential DNA methylation in pediatric acute lymphoblastic leukemia. Genome Biology 14: r105

48. Novershtern N, Subramanian A, Lawton LN, Mak RH, Haining WN, McConkey ME, Habib N, Yosef N, Chang CY, Shay T et al (2011) Densely interconnected transcriptional circuits control cell states in human hematopoiesis. Cell 144: 296–309

49. Panina Y, Germond A, Masui S, Watanabe TM (2018) Validation of Common Housekeeping Genes as Reference for qPCR Gene Expression Analysis During iPS Reprogramming Process. Sci Rep 8: 8716

50. Papaemmanuil E, Rapado I, Li Y, Potter NE, Wedge DC, Tubio J, Alexandrov LB, Van Loo P, Cooke SL, Marshall J et al (2014) RAG-mediated recombination is the predominant driver of oncogenic rearrangement in ETV6-RUNX1 acute lymphoblastic leukemia. Nat Genet 46: 116–125

51. Polak R, Bierings MB, van der Leije CS, Sanders MA, Roovers O, Marchante JRM, Boer JM, Cornelissen JJ, Pieters R, den Boer ML et al (2019) Autophagy inhibition as a potential future targeted therapy for ETV6-RUNX1-driven B-cell precursor acute lymphoblastic leukemia. Haematologica 104: 738–748

52. Popescu D-M, Botting RA, Stephenson E, Green K, Webb S, Jardine L, Calderbank EF, Polanski K, Goh I, Efremova M et al (2019) Decoding human fetal liver haematopoiesis. Nature 574: 365–371

53. Pui CH, Carroll WL, Meshinchi S, Arceci RJ (2011) Biology, risk stratification, and therapy of pediatric acute leukemias: an update. J Clin Oncol 29: 551–565

54. Reimand J, Isserlin R, Voisin V, Kucera M, Tannus-Lopes C, Rostamianfar A, Wadi L, Meyer M, Wong J, Xu C et al (2019) Pathway enrichment analysis and visualization of omics data using g:Profiler, GSEA, Cytoscape and EnrichmentMap. Nat Protoc 14: 482–517

55. Robinson JT, Thorvaldsdottir H, Winckler W, Guttman M, Lander ES, Getz G, Mesirov JP (2011) Integrative genomics viewer. Nat Biotechnol 29: 24–26

56. Schäfer D, Olsen M, Lähnemann D, Stanulla M, Slany R, Schmiegelow K, Borkhardt A, Fischer U (2018) Five percent of healthy newborns have an ETV6-RUNX1 fusion as revealed by DNA-based GIPFEL screening. Blood 131: 821–826

57. Schindler JW, Van Buren D, Foudi A, Krejci O, Qin J, Orkin SH, Hock H (2009) TEL-AML1 Corrupts Hematopoietic Stem Cells to Persist in the Bone Marrow and Initiate Leukemia. Cell Stem Cell 5: 43–53

58. Shannon P, Markiel A, Ozier O, Baliga NS, Wang JT, Ramage D, Amin N, Schwikowski B, Ideker T (2003) Cytoscape: a software environment for integrated models of biomolecular interaction networks. Genome Res 13: 2498–2504

59. Starkova J, Madzo J, Cario G, Kalina T, Ford A, Zaliova M, Hrusak O, Trka J (2007) The Identification of (ETV6)/RUNX1-Regulated Genes in Lymphopoiesis Using Histone Deacetylase Inhibitors in ETV6/RUNX1-Positive Lymphoid Leukemic Cells. Clinical Cancer Research 13: 1726–1735

60. Taylor SC, Nadeau K, Abbasi M, Lachance C, Nguyen M, Fenrich J (2019) The Ultimate qPCR Experiment: Producing Publication Quality, Reproducible Data the First Time. Trends Biotechnol 37: 761–774

61. Teppo S, Laukkanen S, Liuksiala T, Nordlund J, Oittinen M, Teittinen K, Grönroos T, St-Onge P, Sinnett D, Syvänen AC et al (2016) Genome-wide repression of eRNA and target gene loci by the ETV6-RUNX1 fusion in acute leukemia. Genome Res 26: 1468–1477

62. Tirosh I, Izar B, Prakadan SM, Wadsworth MH, Treacy D, Trombetta JJ, Rotem A, Rodman C, Lian C, Murphy G et al (2016) Dissecting the multicellular ecosystem of metastatic melanoma by single-cell RNA-seq. Science 352: 189–196

63. Torrano V, Procter J, Cardus P, Greaves M, Ford AM (2011) ETV6-RUNX1 promotes survival of early B lineage progenitor cells via a dysregulated erythropoietin receptor. Blood 118: 4910–4918

64. Townes FW, Hicks SC, Aryee MJ, Irizarry RA (2019) Feature selection and dimension reduction for single-cell RNA-Seq based on a multinomial model. Genome Biology 20: 295

65. Tsuzuki S, Seto M (2013) TEL (ETV6)-AML1 (RUNX1) Initiates Self-Renewing Fetal Pro-B Cells in Association with a Transcriptional Program Shared with Embryonic Stem Cells in Mice. Stem Cells 31: 236–247

66. Valiron O, Gorka C (1997) Histone H1(0) expression is restricted to progenitor cells during human hematopoiesis. Eur J Cell Biol 72: 39–45

67. van der Weyden L, Giotopoulos G, Rust AG, Matheson LS, van Delft FW, Kong J, Corcoran AE, Greaves MF, Mullighan CG, Huntly BJ et al (2011) Modeling the evolution of ETV6-RUNX1- induced B-cell precursor acute lymphoblastic leukemia in mice. Blood 118: 1041–1051

68. Venugopal B, Baird R, Kristeleit RS, Plummer R, Cowan R, Stewart A, Fourneau N, Hellemans P, Elsayed Y, Mcclue S et al (2013) A Phase I Study of Quisinostat (JNJ-26481585), an Oral Hydroxamate Histone Deacetylase Inhibitor with Evidence of Target Modulation and Antitumor Activity, in Patients with Advanced Solid Tumors. Clinical Cancer Research 19: 4262–4272

69. Vogt M, Dienstbier N, Schliehe-Diecks J, Scharov K, Tu JW, Gebing P, Hogenkamp J, Bilen BS, Furlan S, Picard D et al (2023) Co-targeting HSP90 alpha and CDK7 overcomes resistance against HSP90 inhibitors in BCR-ABL1+ leukemia cells. Cell Death Dis 14: 799

70. Wang L, Hiebert SW (2001) TEL contacts multiple co-repressors and specifically associates with histone deacetylase-3. Oncogene 20: 3716–3725

71. Wiemels JL, Ford AM, Van Wering ER, Postma A, Greaves M (1999) Protracted and variable latency of acute lymphoblastic leukemia after TEL-AML1 gene fusion in utero. Blood 94: 1057–1062

72. Willcockson MA, Healton SE, Weiss CN, Bartholdy BA, Botbol Y, Mishra LN, Sidhwani DS, Wilson TJ, Pinto HB, Maron MI et al (2021) H1 histones control the epigenetic landscape by local chromatin compaction. Nature 589: 293–298

73. Wray JP, Deltcheva EM, Boiers C, Richardson SЕ, Chhetri JB, Brown J, Gagrica S, Guo Y, Illendula A, Martens JHA et al (2022) Regulome analysis in B-acute lymphoblastic leukemia exposes Core Binding Factor addiction as a therapeutic vulnerability. Nature Communications 13: 7124

74. Xing Y, Yu T, Wu YN, Roy M, Kim J, Lee C (2006) An expectation-maximization algorithm for probabilistic reconstructions of full-length isoforms from splice graphs. Nucleic Acids Res 34: 3150–3160

75. Young JC, Wu S, Hansteen G, Du C, Sambucetti L, Remiszewski S, O’Farrell AM, Hill B, Lavau C, Murray LJ (2004) Inhibitors of histone deacetylases promote hematopoietic stem cell self-renewal. Cytotherapy 6: 328–336

76. Zelent A, Greaves M, Enver T (2004) Role of the TEL-AML1 fusion gene in the molecular pathogenesis of childhood acute lymphoblastic leukaemia. Oncogene 23: 4275–4283

